# Sparse latent factor regression models for genome-wide and epigenome-wide association studies

**DOI:** 10.1101/2020.02.07.938381

**Authors:** Basile Jumentier, Kevin Caye, Barbara Heude, Johanna Lepeule, Olivier François

## Abstract

Association of phenotypes or exposures with genomic and epigenomic data faces important statistical challenges. One of these challenges is to account for variation due to unobserved confounding factors, such as individual ancestry or cell-type composition in tissues. This issue can be addressed with penalized latent factor regression models, where penalties are introduced to cope with high dimension in the data. If a relatively small proportion of genomic or epigenomic markers correlate with the variable of interest, sparsity penalties may help to capture the relevant associations, but the improvement over non-sparse approaches has not been fully evaluated yet. Here, we present least-squares algorithms that jointly estimate effect sizes and confounding factors in sparse latent factor regression models. In simulated data, sparse latent factor regression models generally achieved higher statistical performance than other sparse methods, including the least absolute shrinkage and selection operator (LASSO) and a Bayesian sparse linear mixed model (BSLMM). In generative model simulations, statistical performance was slightly lower (while being comparable) to non-sparse methods, but in simulations based on empirical data, sparse latent factor regression models were more robust to departure from the model than the non-sparse approaches. We applied sparse latent factor regression models to a genome-wide association study of a flowering trait for the plant *Arabidopsis thaliana* and to an epigenome-wide association study of smoking status in pregnant women. For both applications, sparse latent factor regression models facilitated the estimation of non-null effect sizes while overcoming multiple testing issues. The results were not only consistent with previous discoveries, but they also pinpointed new genes with functional annotations relevant to each application.

## 1 Introduction

Association studies represent one of the most powerful tool to identify genomic variation correlating with disease states, exposure levels or phenotypes. Those studies are divided into several categories according to the nature of the genomic markers evaluated. For example, genome-wide association studies (GWAS) focus on single-nucleotide polymorphisms in different individuals to estimate disease allele effects (Balding, 2006), while epigenome-wide association studies (EWAS) measure epigenetic marks, such as DNA methylation levels to derive associations between epigenetic variation and exposure levels or to assess effects on phenotypic traits (Rakyan et al., 2011). Despite their success in identifying the genetic architecture of phenotypic traits or genomic targets of exposure, association studies are plagued with the problem of confounding, which arises when unobserved variables correlate with the variable of interest and with the genomic markers simultaneously (Wang et al., 2017). Historical approaches to the confounding issue account for hidden confounders by considering corrections for inflation (Devlin and Roeder, 1999) and empirical null-hypothesis testing methods (Efron, 2004). Alternative approaches evaluate hidden confounders by using linear combinations of observed variables, often called factors. In GWAS, a frequently-used factor approach consists of computing the largest principal components of the genotype matrix, and includes them as covariates in linear regression models (Price et al., 2006). The variable of interest may, however, be collinear to the largest principal components, and removing their effects can result in loss of statistical power. To overcome this issue and increase power, methods based on latent factor regression models have been proposed (Leek and Storey, 2007; Carvalho et al., 2008). Latent factor regression models employ deconvolution methods in which unobserved variables, including batch effects, individual ancestry or tissue cell-type composition are integrated in the regression model by using latent factors. In these models, effect sizes and latent factors are estimated jointly. The latent factor regression framework encompasses several methods which include surrogate variable analysis (SVA, (Leek and Storey, 2007)), latent factor mixed models (LFMM, (Frichot et al., 2013)), residual principal component analysis (Kalaitzis and Lawrence, 2012), direct surrogate variable analysis (Lee et al., 2017), robust reduced-rank regression (She and Chen, 2017) and confounder adjusted testing and estimation (CATE, Wang et al. (2017)). Each method has specific merits relative to some category of association study, and the performances of the methods have been extensively debated in recent surveys (for example, see Kaushal et al. (2017)).

While GWAS and EWAS primarily report significance values, locus-specific effect sizes are summary statistics of crucial importance in interpretation and application of the results (Buniello et al., 2019; Battram et al., 2021). A common property of latent factor regression models is to use regularization parameters inducing constraints on effect size estimates. Among those methods, sparse regression models suppose that a relatively small proportion of all genomic variables correlate with the variable of interest or affect the phenotype, and evaluate associations while avoiding multiple testing problems (Tibshirani, 1996; Hoggart et al., 2008; Wu et al., 2009). Sparse regression models have also been coupled with linear mixed models to combine the benefits of both for polygenic trait studies with Bayesian sparse linear mixed model (BSLMM) (Zhou and Stephens, 2012; Zhou et al., 2013).

In this study, we introduce least-squares algorithms that jointly estimate effect sizes and confounding factors in sparse latent factor regression models. Effect sizes are estimated based on regularized least-squares methods with *L*^1^ and nuclear norm penalties. Thanks to the inclusion of sparsity constraints, the proposed algorithm allows identifying non-null effect sizes without the use of multiple statistical tests. We refer to our approach as *sparse latent factor mixed models* or *sparse* LFMM. We present estimation algorithms for sparse LFMM and theoretical results in the next section. Then we compare the performances of sparse LFMM with other sparse regression models (LASSO, BSLMM) and with non-sparse regression models (SVA, CATE, LFMM with ridge penalty). To illustrate our approach, we used sparse LFMM to perform a GWAS of flowering time for the plant *Arabidopsis thaliana* and to perform an EWAS of smoking status in pregnant women.

## 2 Latent factor regression models

### 2.1 Models

Latent factor regression models evaluate associations between the elements of a response matrix, **Y**, and variables of interest, called *primary* variables, **X**, measured for *n* individuals. The response matrix records *p* markers, which can represent any type of omic data (genotypes, DNA methylation, etc), collected for the individuals. The **X** matrix can also incorporate observed confounders such as age, sex, etc, and its dimension is *n × d*, where *d* represents the total number of variables. Note that latent factor regression models and latent factor mixed models are synonymous terms. Latent factor mixed models contain fixed and latent effects, and those models should not be confused with linear mixed models, which contain fixed and random effects. Latent factor regression models combine fixed and latent effects as follows

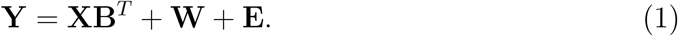

Fixed effect sizes are recorded in the **B** matrix, which has dimension *p × d*. The **E** matrix represents residual errors, and has the same dimension as the response matrix. The matrix **W** is a *latent matrix* of rank *K*, defined by *K* latent factors (Leek and Storey, 2007; Frichot et al., 2013; Wang et al., 2017; Lee et al., 2017). The value of *K* is unknown, and it is generally determined by model choice procedures. The *K* latent factors, **U**, form a matrix of dimension *n × K*, defined from the singular value decomposition of the latent matrix **W** = **UV**^*T*^, where **V** is a semi-orthogonal matrix of dimension *p × K* (Eckart and Young, *1936*). The matrices **U** and **V** are unique up to a change of sign. At the exception of the next paragraph motivating the use of penalized loss functions, this study will, however, not use the decomposition the latent matrix into factor and loading matrices.

Naive statistical estimates for the B and W matrices in equation (1) could be obtained through the minimization of a classical least-squares loss function

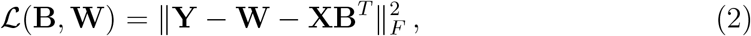

*where* ‖ *·* ‖ _*F*_ is the Frobenius matrix norm. A minimum value of the loss function is attained when **W** is computed as the rank *K* singular value decomposition of **Y** or principal component analysis. In this case, the **B** matrix could be obtained as the estimates of a linear regression of the residual matrix (**Y** *−* **W**) on **X**. To motivate the introduction of regularization terms in the loss function, we remark that the interpretation of the principal components as confounder estimates may be incorrect, because they fail to include any information on the primary variable, and the above construction is problematic. To see it, consider any matrix **P** with dimensions *d × p* and check that

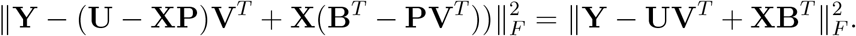

As a consequence, both **B** and (**B** *−* **VP**^*T*^) correspond to valid solutions of the minimization problem, showing there is an infinite space of solutions. To conclude, the loss function needs to be modified in order to warrant dependency of **W** on both **Y** and **X**, and to enable the computation of well-defined solutions.

### 2.2 Sparse estimation algorithms

#### *L*^1^-regularized least-square problem

To solve the problems outlined in the above section, a sparse regularization approach is considered. This approach introduces penalties based on the *L*^1^ norm of the regression coefficients and on the nuclear norm of the latent matrix

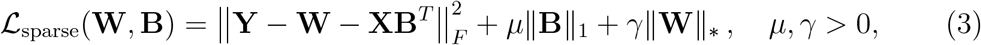

where ‖**B**‖_1_ denotes the *L*^1^ norm of **B**, *µ* is an *L*^1^ regularization parameter, **W** is the latent matrix, ‖**W** ‖_*∗*_ denotes its nuclear norm, and *γ* is a regularization parameter for the nuclear norm. The *L*^1^ penalty induces sparsity on the fixed effects (Tibshirani, 1996), and corresponds to the prior information that not all response variables may be associated with the primary variables. More specifically, the prior implies that a restricted number of rows of the effect size matrix **B** are non-zero. The second regularization term is based on the nuclear norm, and it is introduced to penalize large numbers of latent factors. With these penalty terms, ℒ_sparse_(**W, B**) is a convex function, and convex minimization algorithms can be applied to obtain estimates of **B** and **W** (Mishra et al., 2013).

#### Sparse latent factor mixed model algorithm

Like Tibshirani (1996), let us assume that the primary variables, **X**, are scaled so that **X**^*T*^ **X** = **I** (**I** is the *d × d* identity matrix). Note that the extension to any set of primary variables is not difficult, and that our program implementation is general. We developed a block-coordinate descent method for minimizing the convex loss function ℒ_sparse_(**W, B**) with respect to **B** and **W**. The algorithm is initialized from the null matrix **Ŵ** _0_ = 0, and iterates the following steps.

1. Find 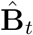 a minimum of the penalized loss function

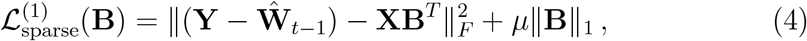
2. Find **Ŵ** _*t*_ a minimum of the penalized loss function

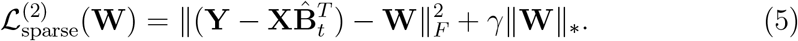

The algorithm cycles through the two steps until a convergence criterion is met or the allocated computing resource is depleted. Each minimization step has a well-defined and unique solution. To see it, note that Step 1 corresponds to an *L*^1^-regularized regression of the residual matrix (**Y** *−* **Ŵ** _*t−* 1_) on the explanatory variables. To compute the regression coefficients, we used the Friedman block-coordinate descent method (Friedman et al., 2007). According to (Tibshirani, 1996), we obtained

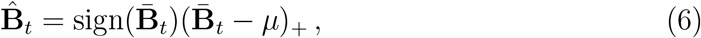

where *s*_+_ = max(0, *s*), sign(*s*) is the sign of *s*, and 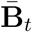 is the linear regression estimate, 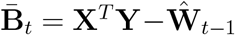. Step 2 consists of finding a low rank approximation of the residual matrix 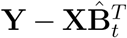 (Cai et al., 2008). This approximation starts with a singular value decomposition (SVD) of the residual matrix, 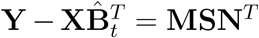, with **M** a unitary matrix of dimension *n × n*, **N** a unitary matrix of dimension *p × p*, and **S** the matrix of singular values (*s*_*j*_)_*j*=1,…,*n*_. Then, we obtain

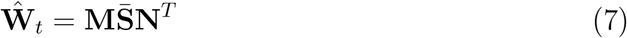

where 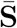 is the diagonal matrix with diagonal terms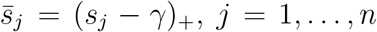. Building on results from (Tseng, 2001), the following statement holds.

##### Theorem 1.

*Let µ >* 0 *and γ >* 0. *Then the block-coordinate descent algorithm cycling through Step 1 and Step 2 converges to estimates of* **W** *and* **B** *defining a global minimum of the penalized loss function ℒ*_sparse_(**W, B**).

Note that the algorithmic complexities of Step 1 and Step 2 are bounded by a term of order *O*(*pn* + *K*(*p* + *n*)). The computing time of sparse LFMM estimates is generally longer than for the CATE algorithm (Wang et al., 2017) or the ridge LFMM algorithm detailed below (Caye et al., 2019). Sparse LFMM needs to perform SVD and projections several times until convergence while CATE and ridge LFMM require a single iteration.

### 2.3 Ridge regression algorithms

Latent factor regression algorithms with ridge penalties were first considered in (Caye et al., 2019). This approach was referred to as ridge LFMM. Since sparse LFMM and ridge LFMM are implemented in the same computer package, we recall how estimates of the parameter matrices **B** and **W** are computed for ridge LFMM. This approach minimizes the following loss function

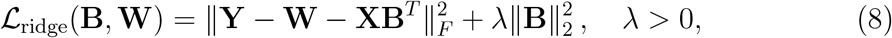

where ‖· ‖_*F*_ is the Frobenius norm, ‖· ‖_2_ is the *L*^2^ norm, and *λ* is a regularization parameter. The minimization algorithm starts with an SVD of the explanatory matrix, **X** = **QΣR**^*T*^, where **Q** is an *n × n* unitary matrix, **R** is an *d × d* unitary matrix and **Σ** is an *n × d* matrix containing the singular values of **X**, denoted by (*σ*_*j*_)_*j*=1,…,*d*_. The ridge LFMM estimates are computed as follows

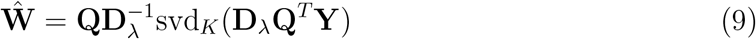

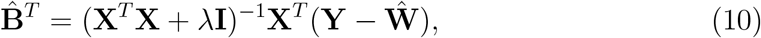

where svd_*K*_(**A**) is the SVD of rank *K* of **A, I** is the *d × d* identity matrix, and **D**_*λ*_ is the *n × n* diagonal matrix with coefficients defined as

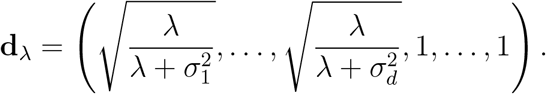

For *λ >* 0, the solution of the regularized least-squares problem is unique (Caye et al., 2019), and the corresponding matrices are called the *ridge estimates*. For completeness, we provide a short proof for this result, stated in (Caye et al., 2019), in the appendix. Using random projections to compute low rank approximations, the complexity of the ridge LFMM algorithm is of order *O*(*n*^2^*p* + *np* log *K*) (Halko et al., 2011). For studies in which the number of samples, *n*, is much smaller than the number of response variables, *p*, computing times of ridge estimates are therefore faster than those of sparse LFMM.

### 2.4 Choice of hyperparameters

#### Number of factors

In order to choose the number of latent factors, *K*, we considered the matrix **D**_*λ*_, defined for the LFMM ridge algorithm, and the unitary matrix **Q**, obtained from an SVD of **X**. The number of latent factors, *K*, was estimated by using spectral analysis of the matrix **D**_0_**Q**^*T*^ **Y**. In real data analyses, we used the scree plot of eigenvalues of the matrix **D**_0_**Q**^*T*^ **Y**, Onalski’s methods based on difference in eigenvalues available in the R package cate (Onatski, 2010), and Leek’s method available in the R package sva (Leek, 2011).

#### Regularization parameters

The *L*^1^-regularization parameter, *µ*, was determined by estimating the proportion of null effect sizes. This proportion was estimated as the proportion of null *p*-values from statistical tests performed with ridge LFMM by using the function pi0est from the package qvalue (Storey et al., 2021). In ridge LFMM, the regularization parameter was set to the default value of *λ* = 10^− 5^. Having set the proportion of non-null effect sizes, *µ* was computed by using the *regularization path* approach proposed by Friedman et al. (2010) as follows. The regularization path algorithm was initialized with the smallest values of *µ* such that

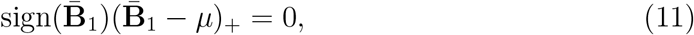

Where 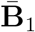 is the linear regression estimate. Then we built a sequence of *µ* values that decreased from the inferred value of the parameter *µ*^max^ to *µ*^min^ = *cµ*^max^. We eventually measured the number of non-null elements in 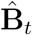, and stopped when the target proportion was reached. The nuclear norm parameter (*γ*) determines the rank of the latent matrix **W**. We used a heuristic approach to evaluate *γ* from the number of latent factors *K*. Based on the singular values (*λ*_1_, …, *λ*_*n*_) of the matrix **D**_0_**Q**^*T*^ **Y**, we set

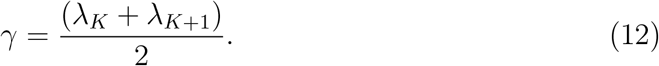

With this value of *γ*, sparse LFMM converged to latent matrix estimates having rank *K* in our experiments.

## 3 Simulation study

In simulation studies, sparse LFMM was compared to two other sparse methods and three non-sparse algorithms, all based on the generative model defined in equation (1). In a first series of model-based experiments, we compared sparse LFMM with three non-sparse approaches, where the performance of each algorithm was measured in scenarios with higher or lower effect sizes and confounding intensities. In a second series with more realistic experiments, we simulated phenotypes based on association with empirical genotypes (Francois and Caye, 2018).

### Estimation algorithms

As a baseline, we used Least Absolute Shrinkage and Selection Operator (LASSO) regression models (Tibshirani, 1996; Friedman et al., 2010). LASSO regression did not include any correction for confounding, and strong biases were expected in effect size estimates. The models were implemented in the R package glmnet, and their regularization parameter was selected by using a 5-fold cross validation approach (Zeng et al., 2017). We also used the Bayesian Sparse Linear Mixed Models (BSLMM) implemented in the GEMMA software (Zhou et al., 2013).

BSLMM is a hybrid method that combines sparse regression models with linear mixed models. BSLMM uses a Markov chain Monte Carlo (MCMC) method to estimate effect sizes. The MCMC burn-in and sampling periods were set to 10,000 (Zeng et al., 2017). To determine the proportion of non-zero effect sizes, two parameters were tuned (*p*_min_ and *p*_max_). Those parameters correspond to the logarithm of the maximum and minimum expected proportions of non-zero effect size. To compare sparse LFMM to non-sparse algorithms, we first used Surrogate Variable Analysis (SVA, (Leek and Storey, 2007)). The algorithm starts with estimating the loading values of a principal component analysis for the residuals of the regression of the response matrix **Y** on **X**. In a second step, SVA determines a subset of response variables exhibiting low correlation with **X**, and uses this subset of variables to estimate the latent factors. SVA was implemented in the R package sva. Next, we implemented the Confounder Adjusted Testing and Estimation (CATE) method (Wang et al., 2017). CATE uses a linear transformation of the response matrix such that the first axis of this transformation is collinear to **X** and the other axes are orthogonal to **X**. CATE was used without negative controls, and it was implemented in the R package cate. We eventually used the ridge LFMM algorithm from the package lfmm (Caye et al., 2019).

### Generative model simulations

In a first series of experiments, we compared sparse LFMM with LASSO and three non-sparse approaches (ridge LFMM, CATE, SVA), and the performance of each algorithm was measured in four scenarios showing high, medium or low effect sizes and confounding intensities. Artificial data were simulated from the generative model defined in equation (1), and the *confounding intensity* was defined as the percentage of variance of the primary variable **X** explained by the latent factors **U**.

More specifically, we simulated a single primary variable (*d* = 1), *K* = 6 latent factors, and a response matrix according to a multivariate Gaussian model (Caye et al., 2019). The joint distribution of (**X, U**) was N(**0, S**), where **S** had diagonal terms 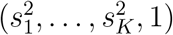, and non-diagonal terms set to zero, except for the covariance between **u**_*k*_ and **X**, which was set to *c*_*k*_*ρ*. The *c*_*k*_ coefficients were sampled from a uniform distribution, and *ρ* was proportional to 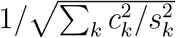. The standard deviations, (*s*_*k*_), were randomly sampled from the interval (2, 6) for *k ≤* 4, and from the interval (0, 1) for *k* = 5, 6. The coefficient of proportionality was chosen so that the confounding intensity could take relatively low (*R*^2^ = 0.1) or high (*R*^2^ = 0.5) values.

To create sparse models, only a small proportion of effect sizes, equal to 0.8%, were allowed to be different from zero. Non-null effect sizes were sampled according to a Gaussian distribution, N(*B*, 0.2), where *B* could take two values, *B* = 0.75 (low value), *B* = 1.5 (medium value) and *B* = 3.0 (high value). Residual errors and loadings, **V**, were sampled according to a standard Gaussian distribution. The dimensions of the response matrix were set to *n* = 400 individuals and *p* = 10, 000 variables. The response matrix was eventually created by simulating from the generative model (equation (1)), and by using a probit link in order to mimic DNA methylation data. Two hundred simulations were performed for each combination of parameters (1200 simulations).

### Results from generative model simulations

Estimation algorithms were used with their default parameters. The number of latent factors was set to *K* = 6 in latent factor models, and the proportion of null-hypothesis was set to 1% in sparse models. Statistical errors (RMSE) of effect size estimates were computed for each method (Figure 1). To provide a reference value for the RMSE, we measured the error made when all effect sizes were estimated as being null (“zero” value or null-model error). The null-model error was equal to 0.069 in lower effect size scenarios and equal to 0.135 in higher effect size scenarios. The RMSEs of sparse LFMM ranged from 0.055 to 0.092, less than those of the null-model. The RMSEs of LASSO were close to the ones of sparse LFMM in low effect size scenarios. In contrast, non-sparse methods led to RMSEs higher than the null-model error, ranging between 0.13 and 0.26 for ridge LFMM and CATE, and rising up to 0.50 for SVA. For the effect sizes associated with causal markers, non-sparse methods reached lower RMSE values than those of sparse methods, ranging between 0.12 and 0.26 for ridge LFMM and CATE, and between 0.60 and 1.03 for sparse LFMM (Figure S1).

**Figure 1.**
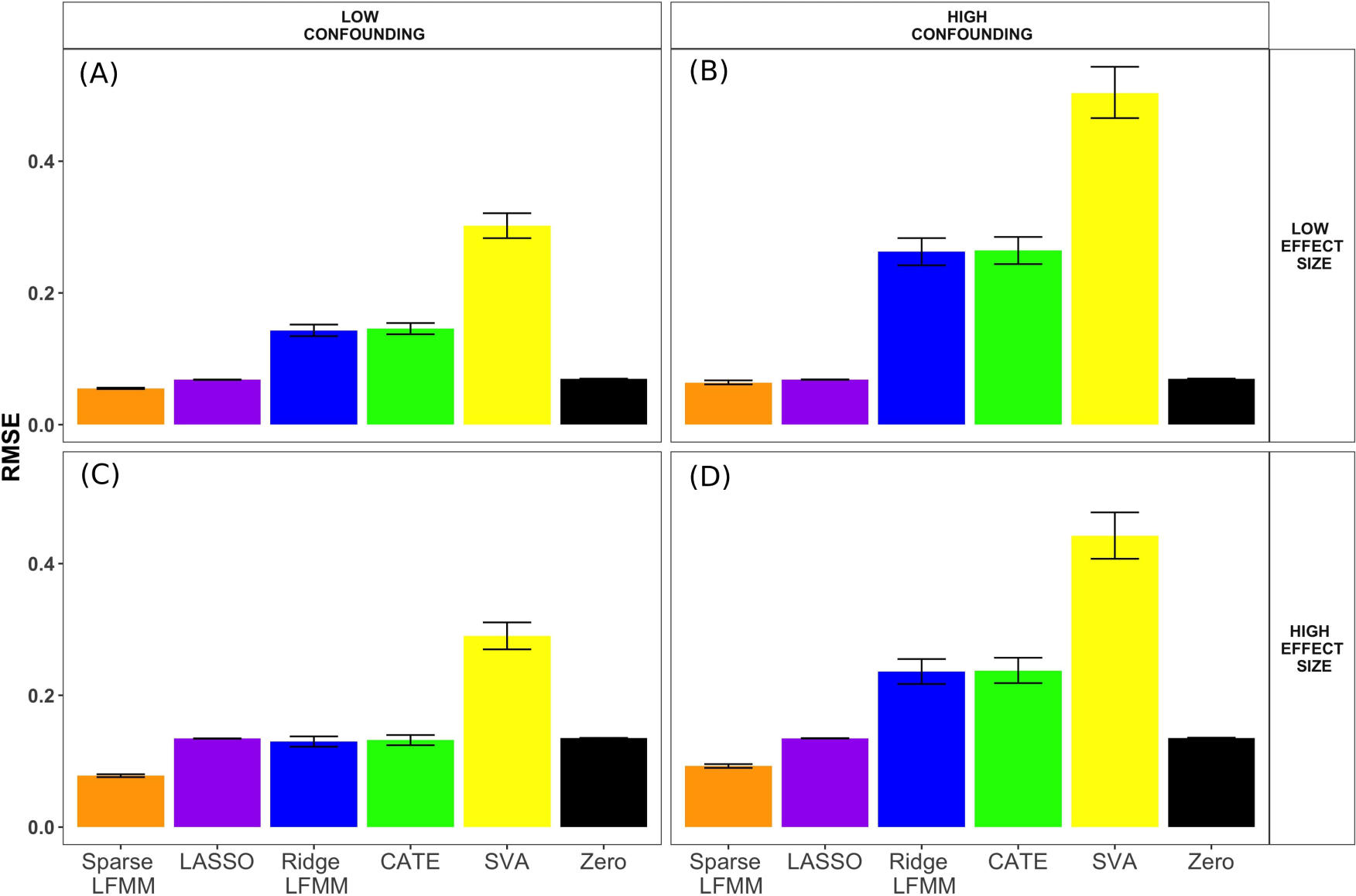
Root Mean Square Error (RMSE) as a function of the effect size of causal markers and confounding intensity. Two sparse methods (sparse LFMM, LASSO) and three non-sparse methods (ridge LFMM, CATE and SVA) were compared. The **zero** value corresponds to an RMSE obtained with all effect sizes set to zero (null-model error). Generative model simulation parameters: (A) Lower effect sizes and confounding intensities (B) Lower effect sizes and higher confounding intensities. (C) Higher effect sizes and lower confounding intensities. (D) Higher effect sizes and confounding intensities.

To evaluate the capabilities of methods to identify true positives, we used precision, which corresponds to the proportion of true positives in a list of positive markers, recall, which is the number of true positives divided by the number of causal markers, and *F* -score, which is the harmonic mean of precision and recall. To compute precision and *F* -score in generative model experiments, a list of *N* = 100 markers with the largest absolute estimated effect sizes was considered for each data set and method. The size of the list, *N*, was chosen as being representative of the number of markers subject to experimental validation. For this value, precision and *F* -score could not be larger than 80% and 89% respectively. To alleviate the choice of a particular value of *N*, we computed precision in (ranked) top hit lists of length *n* for 1 *≤ n ≤ N*, and reported the area under the curve (AUC) normalized by its maximum value. Top hit lists with large AUC values contain more true positives than those with small AUC values.

For all methods, precision and *F* -score were higher in scenarios with higher effect size and lower confounding intensity. In all scenarios, sparse LFMM obtained better scores than SVA. Sparse LFMM reached higher performances than the LASSO when the sizes of the causal effects were medium or high (Figure 2). The situation was reversed only in the case of small effect sizes and for high confounding intensity. In those simulations, sparse LFMM obtained slightly lower scores than ridge LFMM and CATE. The difference was important when the size of causal effects was small (*F* -score *≈* 0.68 against *F* -score *≈* 0.85), but the differences were small for medium and large effect sizes. For large effect sizes, sparse LFMM, ridge LFMM and CATE were close to the maximal *F* -score (0.89) and precision (0.80). AUC scores were strongly correlated to *F* -scores, and the methods were ranked in the same way as with the other performance measures (Figure S2).

**Figure 2.**
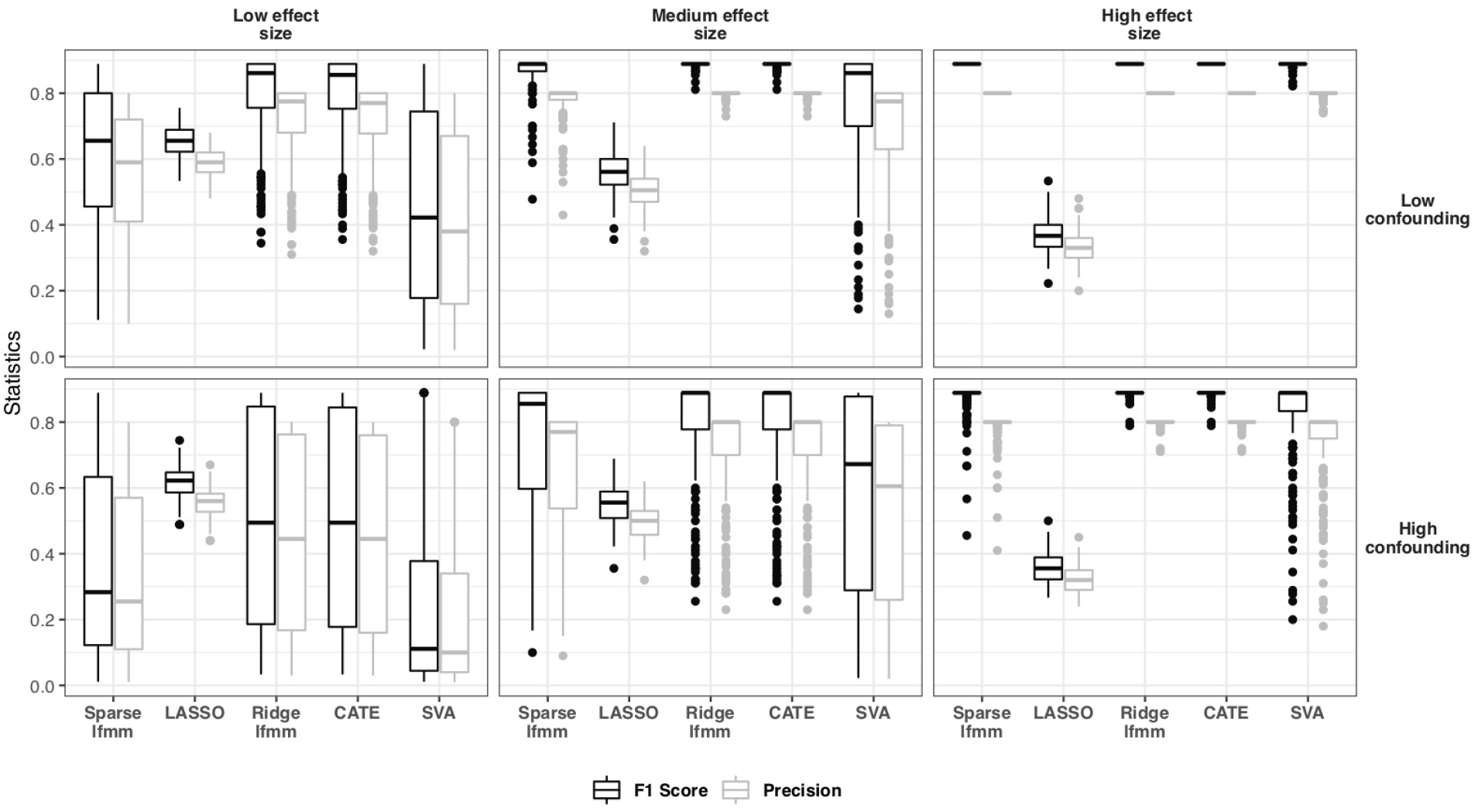
*F*-score and precision for sparse and non-sparse algorithms in generative model simulations. Two sparse methods (sparse LFMM, LASSO) and three non-sparse methods (ridge LFMM, CATE and SVA) were compared for various levels of effect size and confounding intensity in generative model simulations. *F* -score was defined as the harmonic mean of precision and recall.

In summary, sparse LFMM was generally preferable to LASSO and SVA in generative model simulations. Sparse LFMM was associated with the smallest overall statistical error, but the estimates were biased more severely with this method than with non-sparse methods.

### Empirical simulation experiments

In a second series of experiments, we used more realistic simulations to compare sparse LFMM to other sparse and non-sparse methods. Those simulations departed from generative model simulations, and were introduced to evaluate the robustness of effect size estimates in each approach. Phenotypes were simulated for *n* = 162 publicly available Single Nucleotide Polymorphisms (SNPs) genotyped from the fifth chromosome of the model plant *Arabidopsis thaliana* (Atwell et al., 2010). The response matrix contained *p* = 53, 859 SNPs, with minor allele frequency greater than 5%. Phenotypic simulations incorporated realistic features such as gene-by-environment (*G × E*) interactions using environmental variables extracted from a bioclimatic database.

Simulations were performed as follows. Considering a predefined subset of causal markers (*J*), a phenotypic trait, *x*_*i*_, was created for each individual from the following model

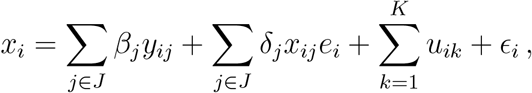

where *y*_*ij*_ represents the genotype of individual *i* at locus *j, e*_*i*_ corresponds to the first principal component of nineteen variables extracted from the bioclimatic database, **u**_*k*_ represents a confounding factor, and *c*_*i*_ is a residual noise. The *β* parameters represent causal variant effect sizes. The number of confounding factors was *K* = 6, and the phenotypes were generated from a combination of five causal SNPs with identical effect sizes. Two values of effect size were used, *β* = 6 (lower effect size) and *β* = 9 (higher effect size). *G × E* interactions were represented by the *δ* parameters. Two values of gene-by-environment interaction were implemented, *G × E* = 0.1 (lower *G × E*) and *G × E* = 0.9 (higher *G × E*). For each parameter combination, two hundred simulations were performed.

### Results from empirical simulation experiments

In empirical simulations, the *F* -score was modified to account for linkage disequilibrium (LD) in the data, and candidate markers within a window of size 10kb around a causal marker were considered to be true discoveries (LD-*r*^2^ *<* 0.2, Francois and Caye (2018)). In lower *G × E* scenarios, sparse LFMM obtained the highest scores (*F* -score in the interval (0.57,0.60), precision in (0.81,0.82), Figure 3AC) compared to BSLMM (*F* -score in (0.36,0.44)), and to non-sparse methods (*F* -score in the interval (0.25, 0.28)). In higher *G × E* scenarios, all methods had poor performances in the lower effect size scenario, but sparse LFMM obtained among the highest *F* -score and precision. When the effect size was higher, sparse LFMM reached higher performances (*F* -score *≈* 0.28 and accuracy *≈* 0.33) than the other methods (Figure 3D). In those realistic simulations, sparse LFMM demonstrated greater robustness to departure from the generative model assumptions than the sparse methods BSLMM and LASSO, and compared favorably to non-sparse methods (ridge LFMM, CATE and SVA).

**Figure 3.**
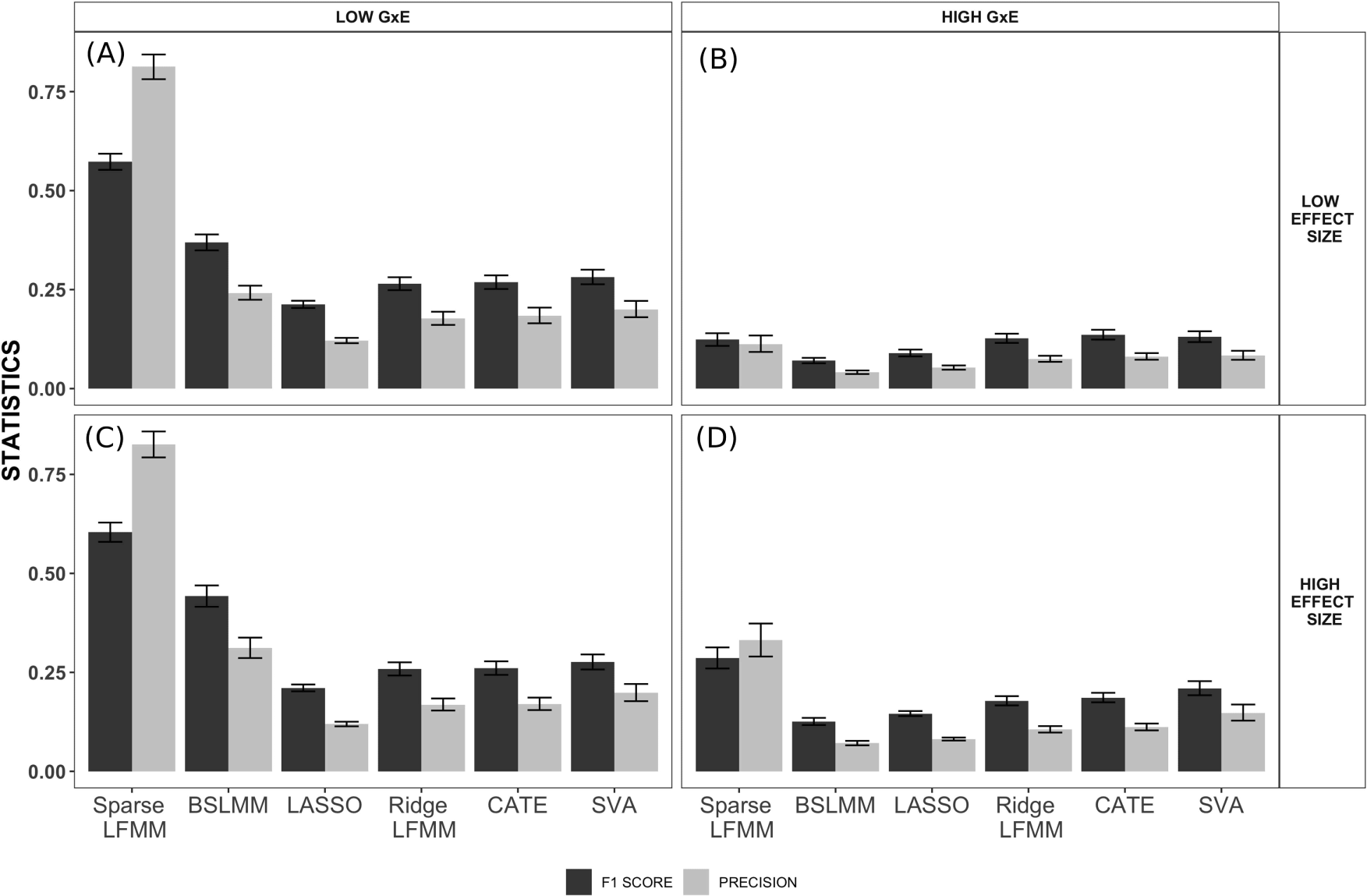
Empirical simulation data (*F*-score and precision). *F* -score and precision as a function of the effect size of the causal markers and of the strength of the interaction between genotype and environment (*G×E*). Three sparse methods (sparse LFMM, BSLMM and LASSO) and three non-sparse methods (ridge LFMM, CATE and SVA) were compared. *F* -score is the harmonic mean of precision and recall. Simulation parameters: (A) Lower effect sizes and lower *G × E* (B) Lower effect sizes and higher *G × E*. (C) Higher effect sizes and lower *G × E*. (D) Higher effect sizes and higher *G × E*.

### Runtimes

Next, we evaluated runtimes for sparse LFMM, and compared those runtimes with BSLMM and ridge LFMM (Figure S3, runtimes measured on an Intel Xeon w-2145 CPU 3.7 GHz). Whatever the number of individuals or markers, ridge LFMM was the fastest method, and sparse LFMM was the slowest method. Higher computation times for sparse LFMM were not surprising because the method iterates many cycles before convergence, whereas ridge LFMM is an exact approach. It took around 2,000 seconds (*≈* 33 minutes) for sparse LFMM to complete runs with *n* = 1, 000 individuals and *p* = 100, 000 markers. While ridge LFMM took a few seconds to compute estimates, around thirty minutes remained an acceptable duration in practical applications. With default values of its MCMC parameters, BSLMM runtimes were of the same order as those of sparse LFMM.

## 4 Data analysis

Sparse LFMM and other methods were used to perform association studies for two distinct types of genomic data including genotypic and epigenetic markers. To illustrate the use of latent factor models in a context where confounding is difficult to control for, we performed a GWAS of flowering time using *p* = 53, 859 SNPs genotyped in chromosome 5 for *n* = 162 European accessions of the model plant *Arabidopsis thaliana*. Our second application evaluated association between smoking during pregnancy and placental DNA methylation for women from a mother-cohort (Heude et al., 2016).

### GWAS of flowering time in *Arabidopsis thaliana*

We considered *n* = 162 European accessions and *p* = 53, 859 SNPs from the fifth chromosome of the *Arabidopsis thaliana* genome to investigate associations with a flowering time phenotype (FT16-TO: 0000344, (Atwell et al., 2010)). FT16 corresponds to the number of days required for an individual plant to reach the flowering stage. In sparse LFMM, the percentage of non-null effect size was set to 1%, in agreement with the proportion of null-hypothesis estimated from non-sparse approaches. The parameters *p*_min_ and *p*_max_ defining sparsity in the BSLMM algorithm were fixed to *p*_min_ = *−* 5 and *p*_max_ = *−* 4 respectively. These values correspond to the logarithm of expected proportions of non-null effect sizes in BSLMM. For all factor methods, the number of latent factors was set to *K* = 10, determined by the eigenvalue methods (see Figure S4).

The sparse methods (sparse LFMM, LASSO, BSLMM) differed in their estimate of the number of null effect sizes (Figure 4ABC, Figure S5). The LASSO approach estimated 99.85% null effect sizes while the proportions were equal to 99.24% and 98.18% for BSLMM and sparse LFMM respectively. The LASSO was the most conservative approach, and sparse LFMM the most liberal one. Sparse LFMM shared 3.9% of hits with LASSO, and 5.5% with BSLMM (Figure S5). Less than 1% of all hits were common to the three approaches. The (non-null) effect sizes for hits varied on distinct scales, with LASSO exhibiting the strongest biases. All sparse methods detected the same top hit at around 4 Mb, corresponding to a SNP located within the *FLC* gene, consistent with the results of (Atwell et al., 2010). The second hit in (Atwell et al., 2010), located in the gene *DOG1*, was also identified by sparse LFMM. BSLMM had more difficulties in identifying previously discovered genes. Given the high correlation – greater than 94 % – between effect sizes obtained with non-sparse methods, we grouped their results by averaging their estimates. Non-sparse methods exhibited effect sizes in a range of values closer to sparse LFMM than to LASSO and BSLMM, but higher statistical errors were observed for those approaches (Figure 4D). Overall, we found a significant correlation between the non-null effect sizes estimated by sparse LFMM and the corresponding effect sizes found by non-sparse methods (*ρ* = 0.8065, *P <* 10^− 16^). In addition, sparse LFMM and the non-sparse methods detected new hits around 13.9 Mb and 6.5 Mb of chr 5, corresponding to the *SAP* and *ACL5* genes respectively.

**Figure 4.**
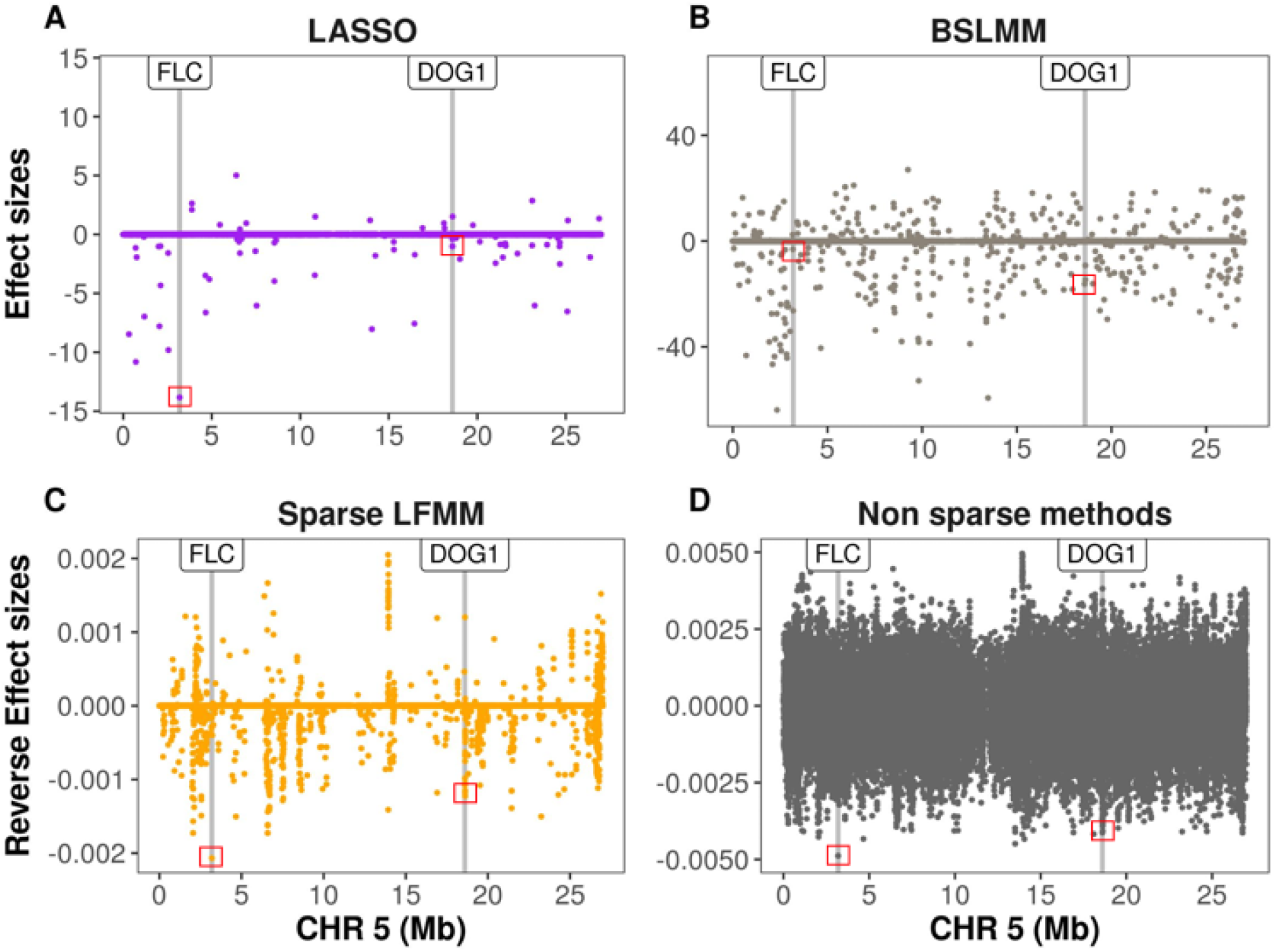
GWAS of flowering time in *A. thaliana* (chromosome 5). A) Effect size estimates for LASSO. B) Effect size estimates for BSLMM. C) Reverse effect size estimates for sparse LFMM. D) Average of reverse effect size estimates for nonsparse methods (ridge LFMM, CATE and SVA). Grey bars represent *Arabidopsis* SNPs associated with the FT16 phenotype in (Atwell et al., 2010), and correspond to the *FLC* and *DOG1* genes.

### EWAS of exposure to smoking during pregnancy

Considering beta-normalized methylation levels at *p* = 425, 878 probed CpG sites for *n* = 668 women, we tested association between smoking status (219 current smoker women and 449 non-current smoker women) and DNA methylation (mDNA) levels in placental tissues. Detailed information on the study population and protocols for placental DNA methylation assessment processing could be found in (Abraham et al., 2018; Rousseaux et al., 2019). For the proportion of non-null effect sizes in sparse LFMM, a conservative value (0.1%) was used (Rousseaux et al., 2019). For latent factor models, the number of latent factors was set to *K* = 7 (see Figure S6).

Using sparse LFMM, the estimated proportion of null effect sizes was equal to 99.698%, for a total number of 1,287 hits (Figure S7). To characterize those 1,287 CpGs, we evaluated whether there was an enrichment of enhancer and promoter regions compared to the methylome (Figure S8 and Figure S9). Promotor and enhancer regions were identified according to the Illumina’s Infinium HumanMethylation450 BeadChip annotations. A fraction of 25.48% were found in enhancer regions, compared to 22.73% for the whole methylome, and 6.83% were found in promoter regions, compared to 19.94% for the whole methylome. We compared the CpGs having the highest effect sizes identified by each method (Figure S10). Sparse LFMM shared 45.3 % of hits with non-sparse models (represented by ridge LFMM), and 2.8 % of hits with LASSO (Table S1). Among the 51 top hits shared by sparse LFMM and ridge LFMM, 25 were found in the body of a gene, 11 were not associated with a gene, 20 were in enhancer regions and 2 in promoter regions. We averaged the effect sizes of non-sparse methods ridge LFMM, CATE and SVA because their correlation was greater than 99%. The results of sparse LFMM agreed with the results of nonsparse methods better than with those of LASSO. The Pearson correlation between the non-null effect sizes estimated by sparse LFMM and the corresponding effect sizes estimated non-sparse methods was equal to *ρ* = 80.38% (*P <* 10^− 16^), whereas the Pearson correlation between non-null effect sizes of sparse LFMM and LASSO was equal to *ρ* = 61.86% (*P <* 10^− 16^). To detail results for a specific genomic region, we focused on chromosome 3, which contained the epigenome-wide top hit for sparse LFMM and for non-sparse methods (cg27402634, located in an enhancer region, Figure 5), and also detected with LASSO (Figure S11). Sparse LFMM hits shared three additional CpGs with non-sparse methods: cg09627057, cg18557837 and cg12662091. Overall, sparse LFMM detected 61 CpGs with non-null effect sizes: 43 were located in genes, 22 in enhancer regions and 6 in promoter regions.

**Figure 5.**
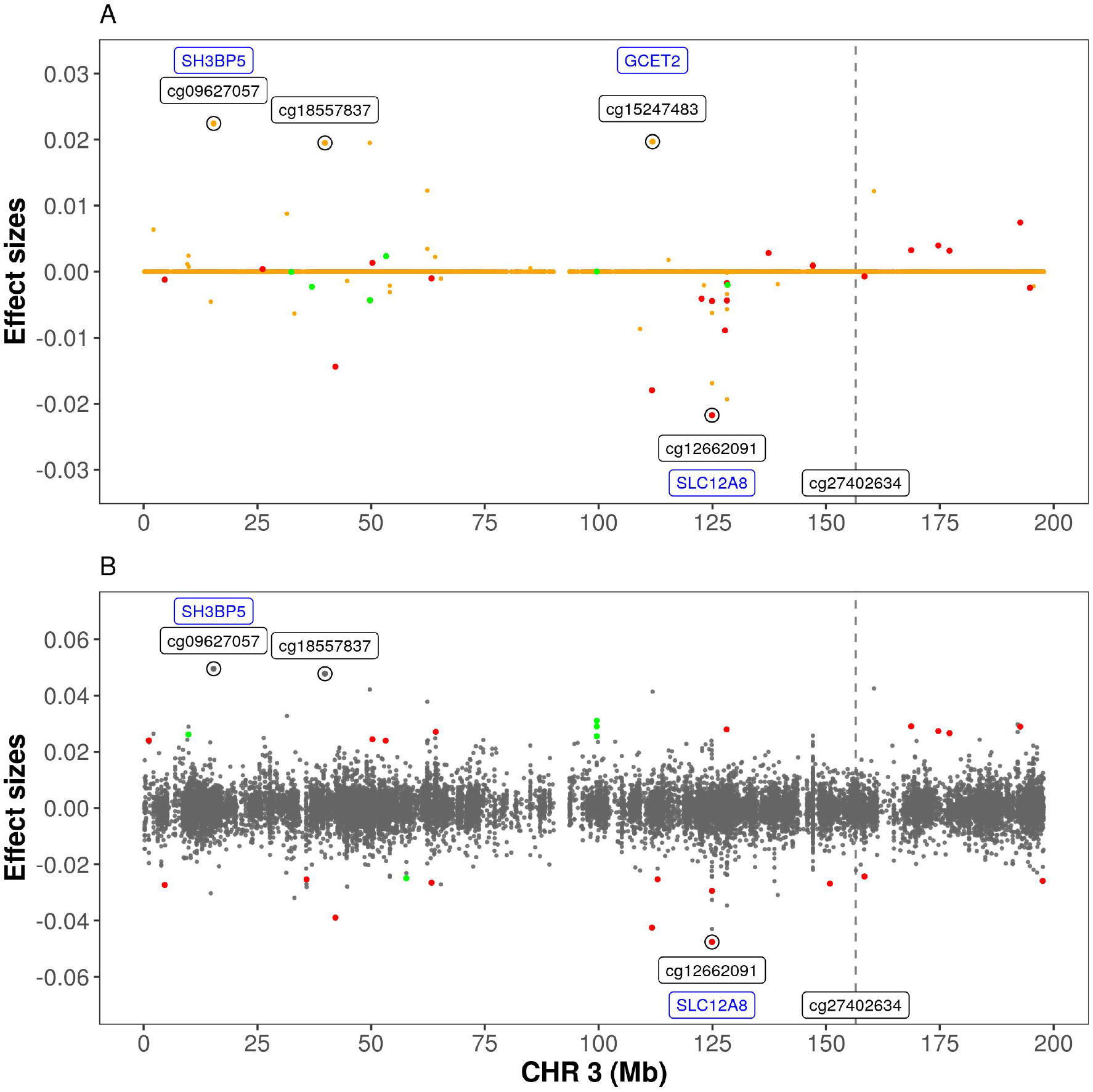
DNA methylation EWAS of smoking status in pregnant women (chromosome 3). A) Estimated effect size for sparse LFMM. The effect size at cg27402634 is equal to *β* = *−* 0.117 (out of range). B) Estimated effect size for nonsparse methods (ridge LFMM, CATE and SVA). The effect size at cg27402634 is equal to *β* = *−* 0.141 (out of range). CpGs with the highest effect sizes are circled (genes are in blue color). Red dots represent CpGs located in enhancer regions. Green dots represent CpGs located in promoter regions.

## 5 Discussion

We introduced sparse latent factor regression methods for the joint estimation of effect sizes and latent factors in genomic and epigenomic association studies. These methods were implemented in the R package lfmm, and are available from the CRAN repository. In generative and in empirical simulations, sparse LFMM obtained higher *F* - score and precision than previously introduced sparse methods, BSLMM and LASSO. Compared to three non-sparse methods (ridge LFMM, CATE and SVA), statistical errors of effect size estimates were overall reduced. In generative model simulations, ridge LFMM and CATE reached higher precision and *F* -score than sparse LFMM. An explanation for this result may be that generative model simulations favor methods for which the estimation of latent factors is unconstrained, and the nuclear norm penalty put on the latent matrix may increase bias in effect size estimates. In contrast, sparse LFMM reached the highest performances in simulations based on empirical data. By being more robust to departure from model assumptions, simulations based on empirical data showed that sparse LFMM can outperform non-sparse methods (see Lotterhos (2019) for a similar conclusion based on population genetic simulations).

A limitation of the sparse LFMM algorithm was its relatively slow execution time, preventing deep exploration of hyperparameter space in real applications. Our strategy was to propose simple guidelines to evaluate the proportion of non-null effect size through an empirical null hypothesis approach, for the number of factors and for the nuclear norm parameter. For causal markers, the effect sizes estimated by sparse LFMM and the corresponding effect sizes estimated by non-sparse methods were strongly correlated. Because effect size estimates had a lower bias in non-sparse methods compared to sparse methods, the results suggest to combine sparse LFMM with a non-sparse method in the following way. At a first stage, sparse LFMM can be used to estimate the support of causal markers having non-null effect sizes. Then ridge LFMM or CATE can be used to estimate the effect sizes of the selected markers. By using sparse LFMM together with a hypothesis testing method, the combined approach alleviates the difficult problem of defining corrections for multiple correlated tests.

In a GWAS of flowering time using 53,859 SNPs in the fifth chromosome of 162 European accessions of the plant *Arabidopsis thaliana*, sparse LFMM identified the *FLC* and *DOG1* genes associated with the FT16 phenotype. The two genes were previously reported as being associated with FT16 in (Atwell et al., 2010). The *FLC* gene plays a central role in flowering induced by vernalization (Sheldon et al., 2000), and *DOG1* is involved in the control of dormancy and seed germination (Nishimura et al., 2018). The second hit of sparse LFMM corresponded to SNPs linked to *SAP*, which is a transcriptional regulator involved in the specification of floral identity (Byzova et al., 1999). This association was also significant for non-sparse methods. In addition, the new method detected SNPs located in the *ACL5* gene, which plays a role in internodal growth and organ size (Hanzawa et al., 1997). In summary, sparse methods facilitated the selection of non-null effect sizes. The results for sparse LFMM were not only consistent with previous discoveries, but they also identified new candidate genes with interesting functional annotations.

Next, we applied sparse LFMM in an EWAS of placental DNA methylation in women exposed to smoking during pregnancy, which is considered an important risk factor for child health (Lumley et al., 2009). The CpG marker cg27402634, which had the highest effect size both in sparse LFMM and in non-sparse methods, was located in an enhancer region, close to the *LEKR1* gene which was associated with birth weight in a GWAS from the Early Growth Genetics (EGG) consortium (http://egg-consortium.org/birth-weight.html). This marker was also detected as a top hit in an independent study of placental methylation and smoking (Morales et al., 2016), and with slightly different study populations and measures of tobacco consumption Rousseaux et al. (2019). In addition to cg27402634, the list of top hits contained several CpGs reported in similar studies, including cg21992501 in the gene *TTC27* (Cardenas et al., 201*9), cg25585967 and cg17823829, respectively in the TRIO* and *KDM5B* genes (Morales et al., 2016; Everson et al., 2019). To characterize the full list of hits, we found an enrichment of enhancer regions and a depletion of promoter regions for CpGs, in agreement with the findings of (Rousseaux et al., 2019). Overall sparse LFMM allowed us to confirm previously discovered associations, and to detect new associations including genes for which methylation changes have detrimental effects on the health of the child.

## Conclusion

Accounting for variation due to unobserved confounding factors is extremely difficult in any type of association study. Assuming that a small proportion of all markers correlate with the exposure or phenotype, we addressed the confounding issue by using sparse latent factor regression models, providing mathematical guarantees that global solutions of least squares estimation problems are proposed. Sparsity constraints in our algorithm allowed the selection of markers alleviating the issue of multiple tests. The application of our method to real data sets highlighted new associations with relevant biological meaning.

### Program Availability

All codes are publicly available, implemented in the R package lfmm available from GitHub for its development version (https://bcm-uga.github.io/lfmm/) and released in the Comprehensive R Archive Network (https://cran.r-project.org/). The scripts used in simulation experiments and in data analyses are available from the following link: https://bcm-uga.github.io/scripts_sagmb.

## Data Availability and Ethics Statement

The *Arabidopsis thaliana* data are publicly available from the 1,001 Genomes database (https://1001genomes.org/). The EDEN individual-level data have restricted access owing to ethical and legal conditions in France. They are available upon request from the EDEN steering committee at and through collaborations with the principal investigators of EDEN. The EDEN cohort received approval from the ethics committee (CCPPRB) of Kremlin Bicêtre and from the French data privacy institution “Commission Nationale de l’Informatique et des Libertés (CNIL)”. Written consent was obtained from the mother for herself and for the offspring.

## Mathematical proofs

This section provides mathematical proofs for the results stated in section 2.

### Theorem 1.

*Let µ >* 0 *and γ >* 0. *Then the block-coordinate descent algorithm cycling through Step 1 and Step 2 converges to estimates of* **W** *and* **B** *defining a global minimum of the penalized loss function* ‖_sparse_(**W, B**).

*Proof*. The proof arguments are based on a result of (Tseng, 2001). Consider the Cartesian product of closed convex sets *A* = *A*_1_ *× A*_2_ *× … × A*_*m*_, and let *f* (**z**) be a continuous convex function defined on *A* and such that

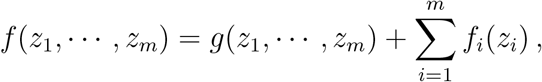

where *g*(**z**) is a differentiable convex function, and for each *i* = 1, …, *m, f*_*i*_(*z*_*i*_) is a continuous convex function. Let (**z**^*t*^) be the sequence of values defined by the following block-coordinate descent algorithm

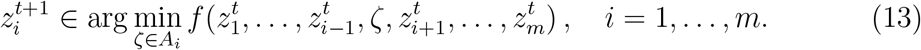

Then a limit point of the sequence (**z**^*t*^) defines a global minimum of the function *f* (**z**). The theorem’s proof is a consequence of the convexity of the penalized loss function ‖_sparse_(**W, B**), and the fact that we can write

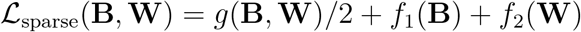

where 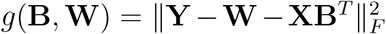 is a differentiable convex function, and 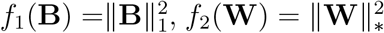 are continuous convex functions. Tseng’s result can be applied with the function *f* (**B, W**) = *ℒ*_sparse_(**B, W**) to conclude the proof (see also (Bertsekas, 1995)).

### Theorem 2.

*Let λ >* 0 *and assume* 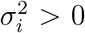 *for all i* = 1, …, *d. The estimates* **Ŵ** *and* 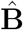 *computed as follows*

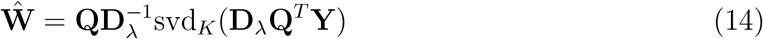

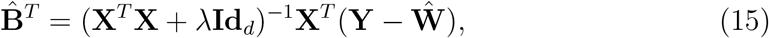

*where* svd_*K*_(**A**) *is the rank K SVD of the matrix* **A, Id**_*d*_ *is the d × d identity matrix, and* **D**_*λ*_ *is the n × n diagonal matrix with coefficients defined as*

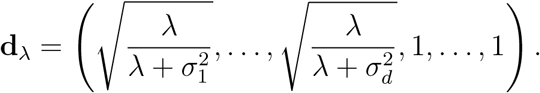

*define a global minimum of the penalized loss function ℒ*_ridge_(**B, W**).

*Proof*. Given **W**, a global minimum for *ℒ*_ridge_(**B, W**) is obtained with the ridge estimates for a linear regression of the response matrix **Y** *−* **W** on **X**.

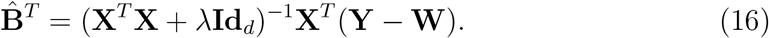

Thus, the problem amounts to minimizing the function 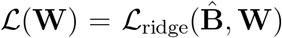 with respect to **W**. By definition of the **D**_*λ*_ and **Q** matrices, the loss function rewrites as

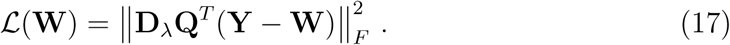

Minimizing the above loss function is equivalent to finding the best approximation of rank *K* for the matrix **D**_*λ*_**Q**^*T*^ **Y**. According to (Eckart and Young, 1936), this approximation is given by the rank *K* singular value decomposition of **D**_*λ*_**Q**^*T*^ **Y**. Eventually we obtain that

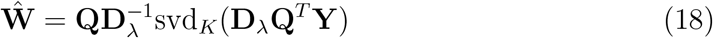

defines the unique global minimum of the *ℒ* (**W**) function.

## Acknowledgments

This article was developed in the framework of the Grenoble Alpes Data Institute, supported by the French National Research Agency under the Investissements d’Avenir program (ANR-15-IDEX-02). It received support from LabEx PERSY-VAL Lab, ANR-11-LABX-0025-01, and from the French National Research Agency (Agence Nationale pour la Recherche) ETAPE, ANR-18-CE36-0005. The EDEN mother-child study was supported by Foundation for medical research (FRM), National Agency for Research (ANR), National Institute for Research in Public health (IRESP), French Ministry of Health (DGS), French Ministry of Research, INSERM Bone and Joint Diseases National Research (PRO-A), and Human Nutrition National Research Programs, Nestlé, French National Institute for Population Health Surveillance (InVS), French National Institute for Health Education (INPES), the European Union FP7 programmes (FP7/2007-2013, HELIX, ESCAPE, ENRIECO, Medall projects), Diabetes National Research Program, French Agency for Environmental Health Safety (ANSES), Mutuelle Générale de l’Education Nationale (MGEN), French national agency for food security, French-speaking association for the study of diabetes and metabolism (ALFEDIAM). We thank all the participants and members of the EDEN mother-child cohort study group.

