## Supplementary materials for "Sparse latent factor regression models for genome-wide and epigenome-wide association studies"

**Table S1.** EWAS of smoking status in pregnant women. List of fifty-one top hits shared between sparse and ridge LFMM analyses.

| CpG | Chr | Position (built 37) | Gene | Location in gene | Location of CpG | Enhancer | Mean methylation level | SE | Beta Sparse LFMM | BETA Ridge LFMM | Rank Sparse LFMM | Rank Ridge LFMM | Beta LASSO |
|---|---|---|---|---|---|---|---|---|---|---|---|---|---|
| **cg27402634** | 3 | 156536860 | | | S_Shore | Yes | 0,64951 | 0,00421 | -0,11735 | -0,14040 | 1 | 1 | -2,14621 |
| **cg27467876** | 8 | 22266134 | SLC39A14 | Body | | No | 0,61827 | 0,01312 | -0,04729 | -0,07197 | 2 | 3 | 0,00000 |
| **cg04998327** | 11 | 103453960 | | | | Yes | 0,52602 | 0,01113 | -0,04674 | -0,07681 | 3 | 2 | 0,00000 |
| **cg03449867** | 15 | 28200653 | OCA2 | Body | | No | 0,45392 | 0,01087 | 0,03852 | 0,06030 | 4 | 13 | 0,00000 |
| **cg08629394** | 12 | 96188933 | | | S_Shelf | No | 0,56171 | 0,00881 | 0,03775 | 0,05776 | 5 | 16 | 0,00000 |
| **cg23698271** | 10 | 121346762 | TIAL1 | Body | | Yes | 0,74335 | 0,00970 | -0,03470 | -0,06649 | 6 | 6 | 0,00000 |
| **cg21211688** | 9 | 136403935 | ADAMTSL2 | Body | S_Shelf | No | 0,62514 | 0,01225 | 0,03447 | 0,06873 | 7 | 4 | 0,00000 |
| **cg21463262** | 13 | 113539522 | ATP11A | 3'UTR | N_Shore | Yes | 0,47291 | 0,01243 | -0,03447 | -0,06176 | 8 | 11 | 0,00000 |
| **cg00033213** | 8 | 144399335 | TOP1MT | Body | Island | No | 0,49779 | 0,01270 | -0,03432 | -0,06546 | 9 | 8 | 0,00000 |
| **cg09785377** | 15 | 60644157 | ANXA2 | Body | | Yes | 0,65664 | 0,01083 | 0,03431 | 0,05556 | 10 | 18 | 0,00000 |
| **cg03796003** | 16 | 2746544 | KCTD5 | Body | Island | No | 0,58449 | 0,01240 | -0,03379 | -0,06557 | 11 | 7 | 0,00000 |
| **cg04835489** | 1 | 52624196 | ZFYVE9 | 5'UTR | | Yes | 0,70411 | 0,00341 | 0,03353 | 0,06427 | 12 | 9 | 0,89825 |
| **cg21566433** | 15 | 101936156 | PCSK6 | Body | N_Shelf | No | 0,63087 | 0,01116 | -0,03338 | -0,06792 | 13 | 5 | 0,00000 |
| **cg21992501** | 2 | 32872637 | TTC27 | Body | | Yes | 0,66651 | 0,00359 | 0,03072 | 0,05696 | 14 | 17 | 0,33409 |
| **cg19949776** | 15 | 51236304 | LOC100132724 | TSS200 | | No | 0,57014 | 0,01232 | 0,03005 | 0,06198 | 15 | 10 | 0,00000 |
| **cg12150991** | 11 | 9113146 | SCUBE2 | 1stExon | Island | No | 0,29875 | 0,00913 | -0,02771 | -0,05450 | 16 | 19 | 0,00000 |
| **cg17823829** | 1 | 202765754 | KDM5B | Body | | Yes | 0,67778 | 0,00354 | 0,02756 | 0,05857 | 17 | 15 | 0,20159 |
| **cg07874011** | 7 | 3134592 | | | | No | 0,42951 | 0,00943 | 0,02701 | 0,04920 | 18 | 35 | 0,00000 |
| **cg20550012** | 7 | 157370486 | PTPRN2 | Body | S_Shore | No | 0,53862 | 0,00955 | 0,02663 | 0,05320 | 19 | 21 | 0,00000 |
| **cg03216697** | 6 | 31239295 | HLA-C | Body | Island | No | 0,28611 | 0,01118 | 0,02514 | 0,05015 | 21 | 29 | 0,00000 |
| **cg24642483** | 1 | 159261560 | FCER1A | 5'UTR | | No | 0,46717 | 0,00695 | 0,02500 | 0,05859 | 23 | 14 | 0,01213 |
| **cg25585967** | 5 | 14452105 | TRIO | Body | | Yes | 0,71265 | 0,00291 | 0,02478 | 0,05252 | 24 | 22 | 0,87552 |
| **cg07755718** | 8 | 8639450 | | | | No | 0,77787 | 0,00710 | -0,02404 | -0,04481 | 27 | 60 | 0,00000 |
| **cg26089705** | 20 | 43371550 | | | N_Shelf | No | 0,38208 | 0,00775 | -0,02379 | -0,05130 | 28 | 25 | 0,00000 |
| **cg21472506** | 2 | 63283967 | OTX1 | 3'UTR | Island | Yes | 0,50118 | 0,00830 | -0,02369 | -0,04835 | 29 | 39 | 0,00000 |
| **cg05161773** | 17 | 75378036 | SEPT9 | 5'UTR | | Yes | 0,73301 | 0,01117 | 0,02327 | 0,04818 | 30 | 42 | 0,00000 |
| **cg24844518** | 5 | 156811669 | CYFIP2 | Body | | Yes | 0,60704 | 0,01094 | -0,02307 | -0,06047 | 31 | 12 | 0,00000 |
| **cg01163842** | 14 | 95235125 | GSC | Body | Island | Yes | 0,26699 | 0,00932 | -0,02269 | -0,04501 | 32 | 58 | 0,00000 |
| **cg07703391** | 1 | 40226045 | BMP8B | 3'UTR | | No | 0,66945 | 0,01145 | -0,02250 | -0,04831 | 33 | 40 | 0,00000 |
| **cg09627057** | 3 | 15377670 | SH3BP5 | 5'UTR | S_Shelf | No | 0,96734 | 0,00046 | 0,02245 | 0,04932 | 34 | 33 | 0,00000 |
| **cg23853165** | 10 | 42864912 | LOC441666 | TSS1500 | S_Shore | No | 0,59014 | 0,00732 | 0,02234 | 0,05378 | 36 | 20 | 0,01383 |
| **cg23548640** | 8 | 789977 | | | Island | No | 0,74132 | 0,00690 | 0,02233 | 0,04820 | 37 | 41 | 0,00159 |
| **cg25178500** | 12 | 88827943 | | | | Yes | 0,91084 | 0,00125 | -0,02207 | -0,04766 | 38 | 46 | 0,00000 |
| **cg17149911** | 2 | 69812277 | AAK1 | Body | | Yes | 0,64968 | 0,01112 | 0,02187 | 0,05221 | 39 | 23 | 0,00000 |
| **cg12662091** | 3 | 124931535 | SLC12A8 | 5'UTR | Island | Yes | 0,26770 | 0,00668 | -0,02174 | -0,04782 | 40 | 44 | 0,00000 |
| **cg27586797** | 5 | 13664584 | | | | Yes | 0,46900 | 0,01166 | 0,02159 | 0,04703 | 41 | 51 | 0,00000 |
| **cg23162598** | 7 | 157345746 | PTPRN2 | Body | N_Shore | No | 0,38177 | 0,00930 | -0,02108 | -0,04783 | 44 | 43 | 0,00000 |
| **cg15790037** | 17 | 37321490 | ARL5C | Body | Island | No | 0,23476 | 0,00799 | -0,02107 | -0,04883 | 45 | 37 | 0,00000 |
| **cg19300401** | 6 | 16962712 | | | | Yes | 0,66063 | 0,01184 | -0,02087 | -0,05096 | 46 | 26 | 0,00000 |
| **cg07480176** | 7 | 94144926 | CASD1 | Body | | Yes | 0,48100 | 0,00882 | 0,02084 | 0,04425 | 47 | 64 | 0,00000 |
| **cg27119456** | 10 | 42863173 | LOC441666 | Body | Island | No | 0,37859 | 0,00703 | -0,02082 | -0,05067 | 48 | 27 | 0,00000 |
| **cg22280068** | 11 | 285037 | NLRP6 | Body | N_Shelf | No | 0,68526 | 0,00898 | -0,02081 | -0,04991 | 49 | 31 | 0,00000 |
| **cg07938743** | 2 | 63283939 | OTX1 | 3'UTR | Island | No | 0,60780 | 0,00773 | -0,02062 | -0,04739 | 50 | 49 | 0,00000 |
| **cg18662228** | 2 | 236867804 | AGAP1 | Body | Island | No | 0,48277 | 0,01185 | -0,02057 | -0,04917 | 51 | 36 | 0,00000 |
| **cg17830140** | 19 | 626119 | POLRMT | Body | S_Shelf | No | 0,82585 | 0,00773 | -0,02040 | -0,04669 | 52 | 54 | 0,00000 |
| **cg00345083** | 1 | 4725584 | AJAP1 | Body | N_Shore | No | 0,50177 | 0,01122 | 0,02021 | 0,04705 | 53 | 50 | 0,00000 |
| **cg00868875** | 18 | 24127237 | KCTD1 | 5'UTR | Island | No | 0,77468 | 0,00475 | -0,02019 | -0,04465 | 54 | 62 | -0,05191 |
| **cg05593887** | 7 | 77827379 | MAGI2 | Body | | Yes | 0,56928 | 0,01178 | 0,01971 | 0,04941 | 56 | 32 | 0,00000 |
| **cg18557837** | 3 | 39847605 | | | N_Shelf | No | 0,19914 | 0,00886 | 0,01951 | 0,04759 | 59 | 47 | 0,00000 |
| **cg24527560** | 10 | 42863508 | LOC441666 | TSS200 | S_Shore | No | 0,29778 | 0,00712 | -0,01917 | -0,04769 | 63 | 45 | 0,00000 |
| **cg07271471** | 13 | 24788196 | SPATA13 | 5'UTR | | No | 0,69031 | 0,01055 | 0,01907 | 0,04496 | 65 | 59 | 0,00000 |

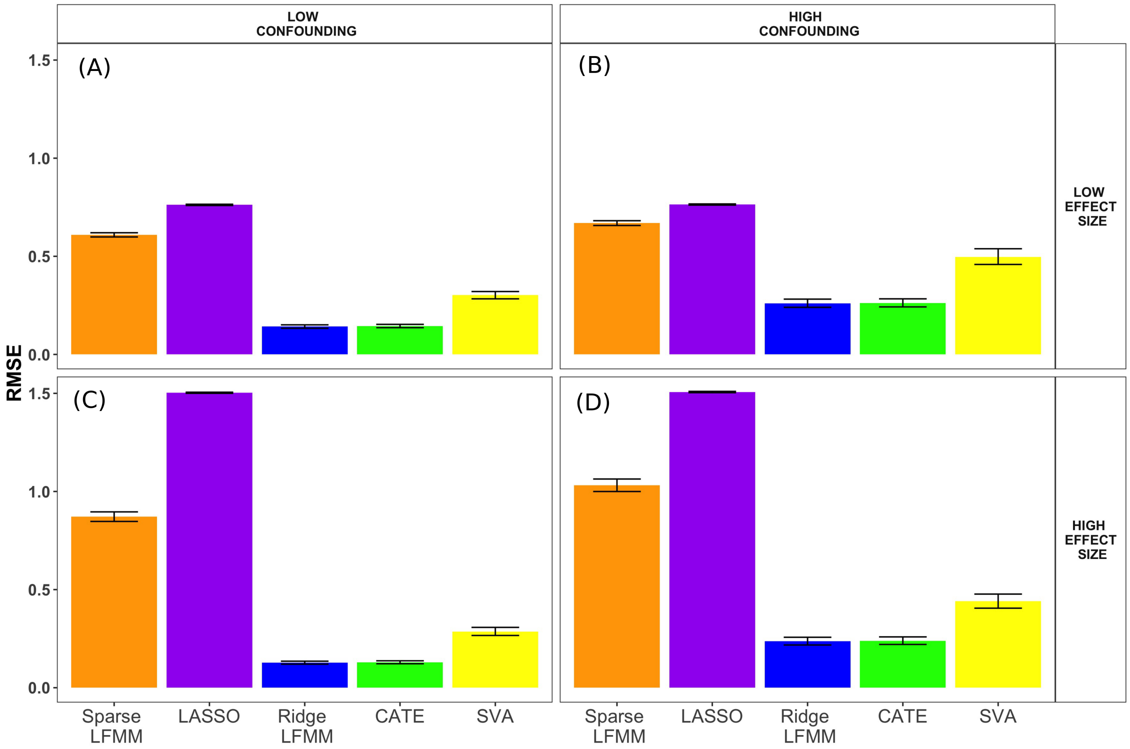


**Figure S1. Generative model simulations (RMSE for causal markers only).** Root Mean Square Error (RMSE) of causal effect sizes as a function of the effect size of the causal markers and of the confounding intensity. Two sparse methods (sparse LFMM, LASSO) and three non-sparse methods (ridge LFMM, CATE and SVA) were compared. Simulation parameters: (A) Lower effect sizes and confounding intensities (B) Lower effect sizes and higher confounding intensities. (C) Higher effect sizes and lower confounding intensities. (D) Higher effect sizes and confounding intensities.

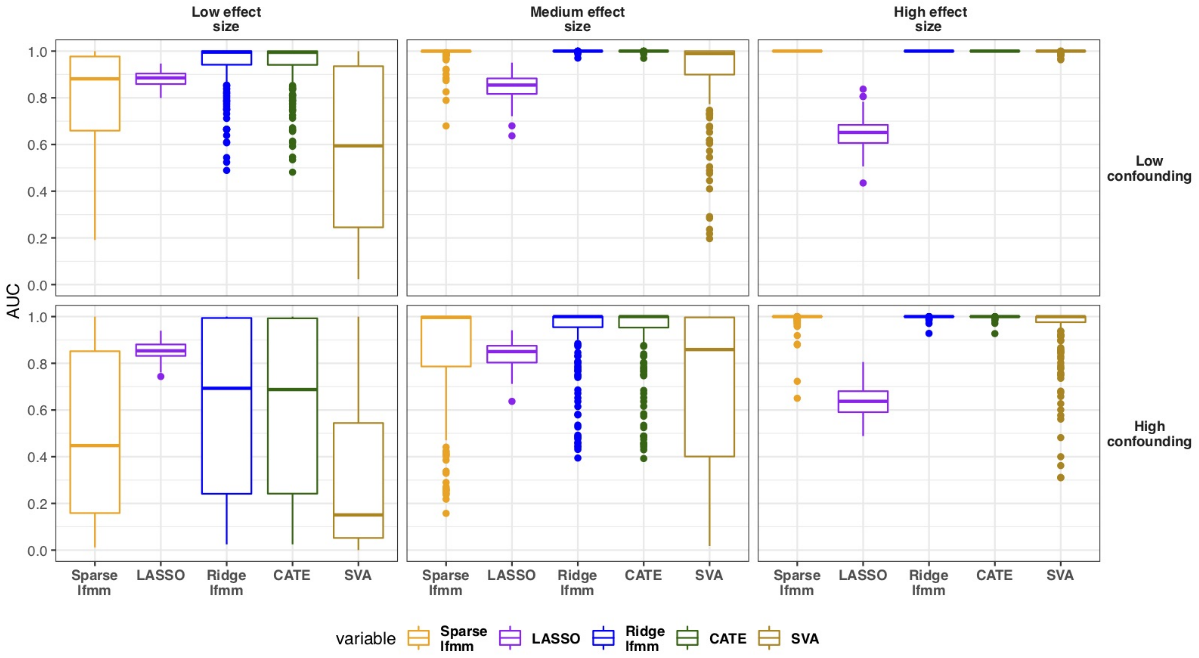


**Figure S2. AUC for sparse and non-sparse algorithms in generative model simulations.** Two sparse methods (sparse LFMM, LASSO) and three non-sparse methods (ridge LFMM, CATE and SVA) were compared. AUC was computed as the normalized area under the curve of precision for the $N$ top hits ($1 \leq N \leq 100$, $p = 10,000$ markers, 80 causal markers). See main text for values of effect sizes and confounding intensities in generative model simulations.

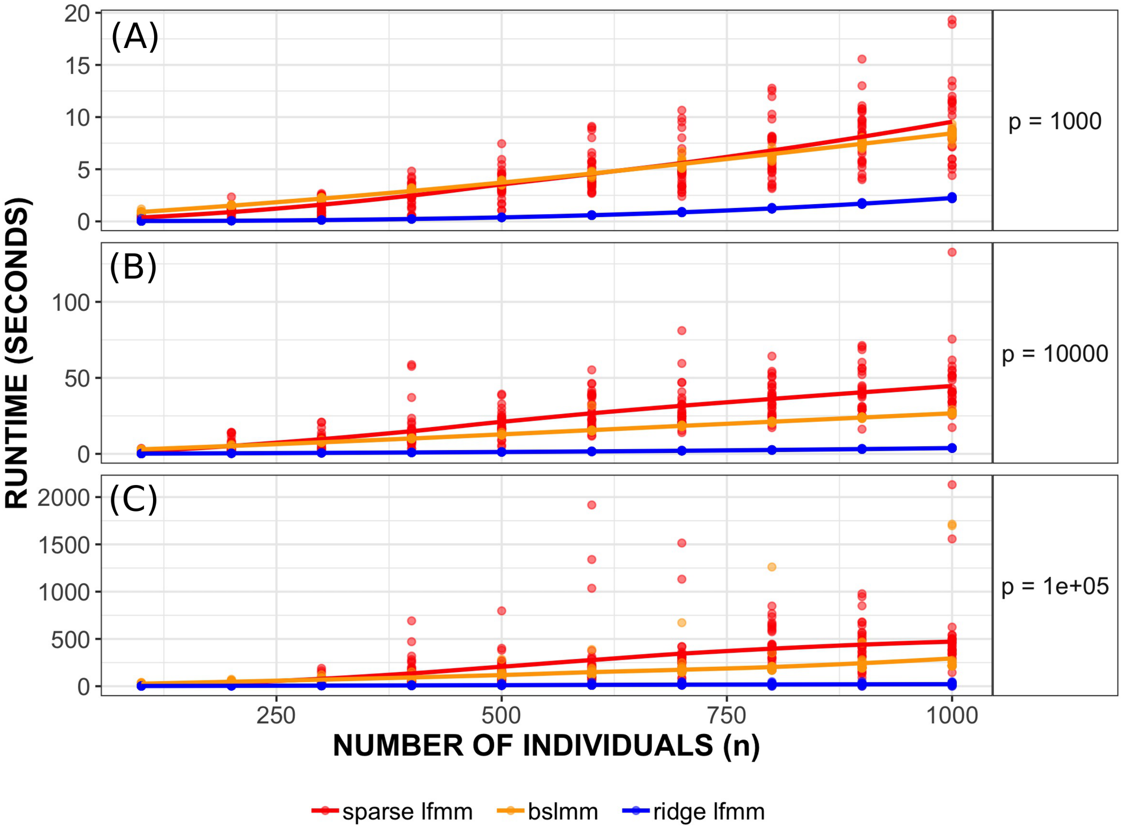


**Figure S3. Comparison of runtimes of three methods.** Runtimes as a function of the number of markers ($p$) and the number of individuals ($n$). (A) $p = 1000$. (B) $p = 10,000$. (C) $p = 100,000$.

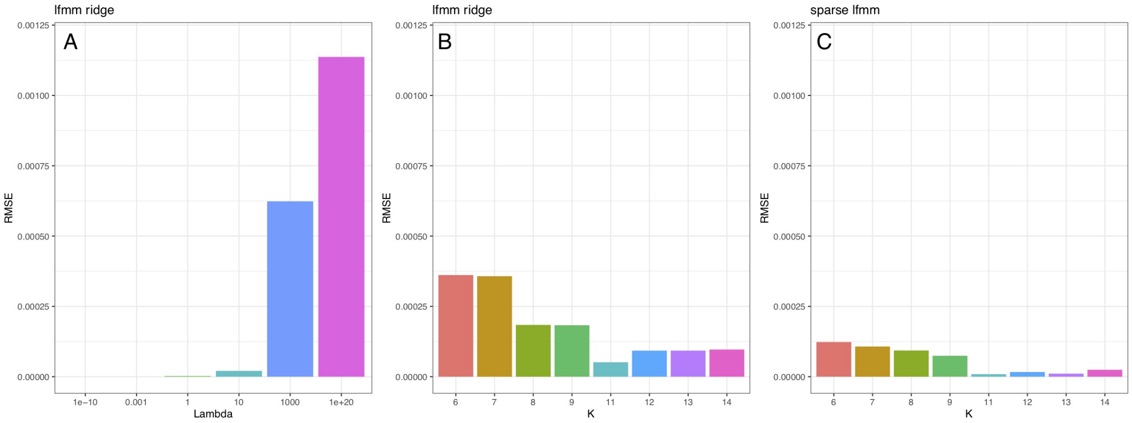


**Figure S4. Impact of hyperparameters in a GWAS of a flowering trait.** Effect sizes were estimated with ridge LFMM for several values of hyperparameters $\lambda$ and $K$ (number of factors), and for sparse LFMM with several values of $K$. RMSE were calculated with respect to the effect sizes reported in the main text, obtained with $\lambda = 10^{-5}$ and $K = 10$. A) Ridge LFMM with ridge parameter $\lambda$, B) Ridge LFMM with number of factors, $K$. C) Sparse LFMM with number of factors, $K$. The results were similar for a large range of values of $\lambda$ and $K$.

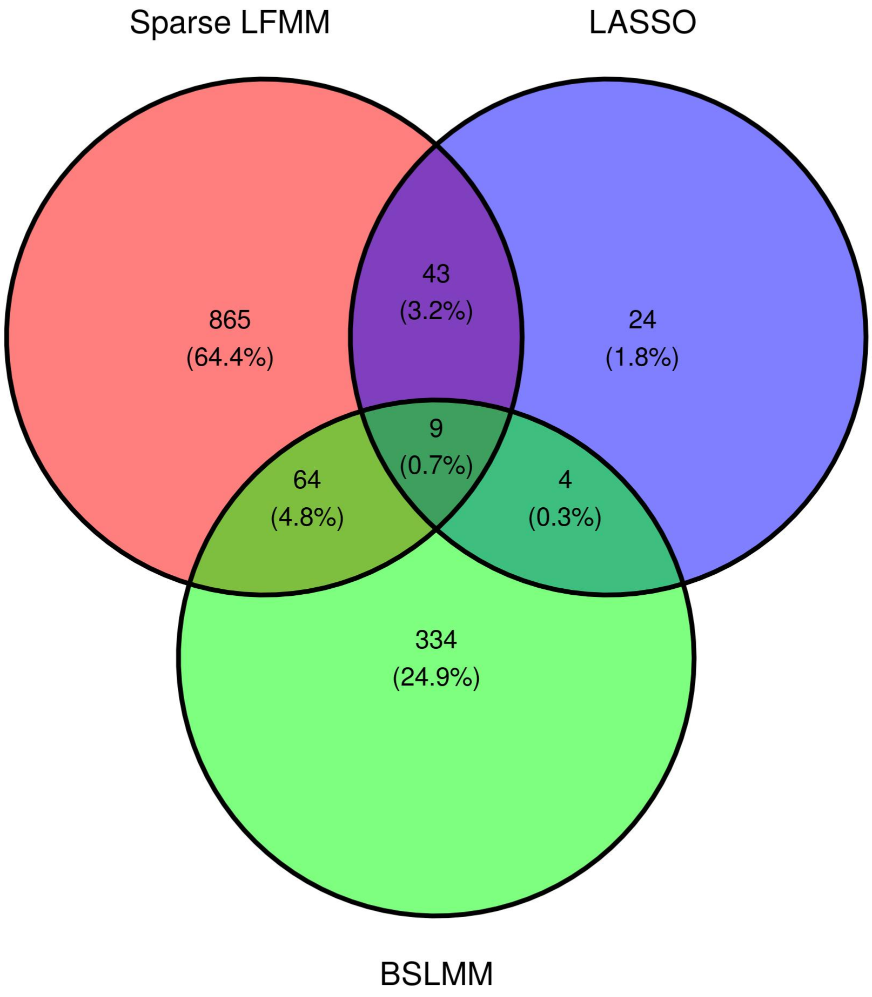


**Figure S5. GWAS of a flowering trait with sparse LFMM, LASSO and BSLMM.** Venn diagram of SNP hits associated with the FT16 phenotype in each approach. The hits correspond to SNPs having non-null effect size estimates.

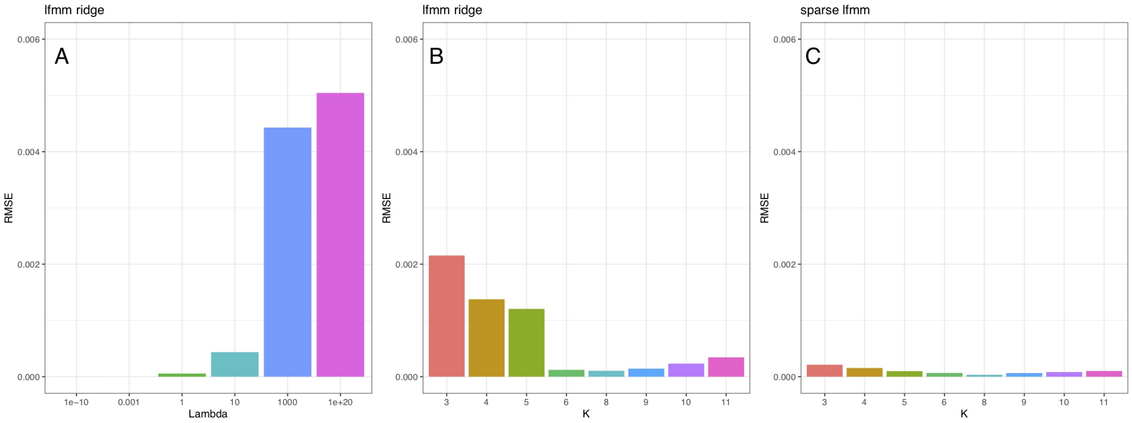


**Figure S6. Impact of hyperparameters in an EWAS of smoking status in pregnant women.** Effect sizes were estimated with ridge LFMM for several values of hyperparameters $\lambda$ and $K$ (number of factors), and for sparse LFMM with several values of $K$. RMSE were calculated with respect to the effect sizes reported in the main text, obtained with $\lambda = 10^{-5}$ and $K = 7$. A) Ridge LFMM with ridge parameter $\lambda$, B) Ridge LFMM with number of factors, $K$. C) Sparse LFMM with number of factors, $K$. The results were similar for a large range of values of $\lambda$ and $K$.

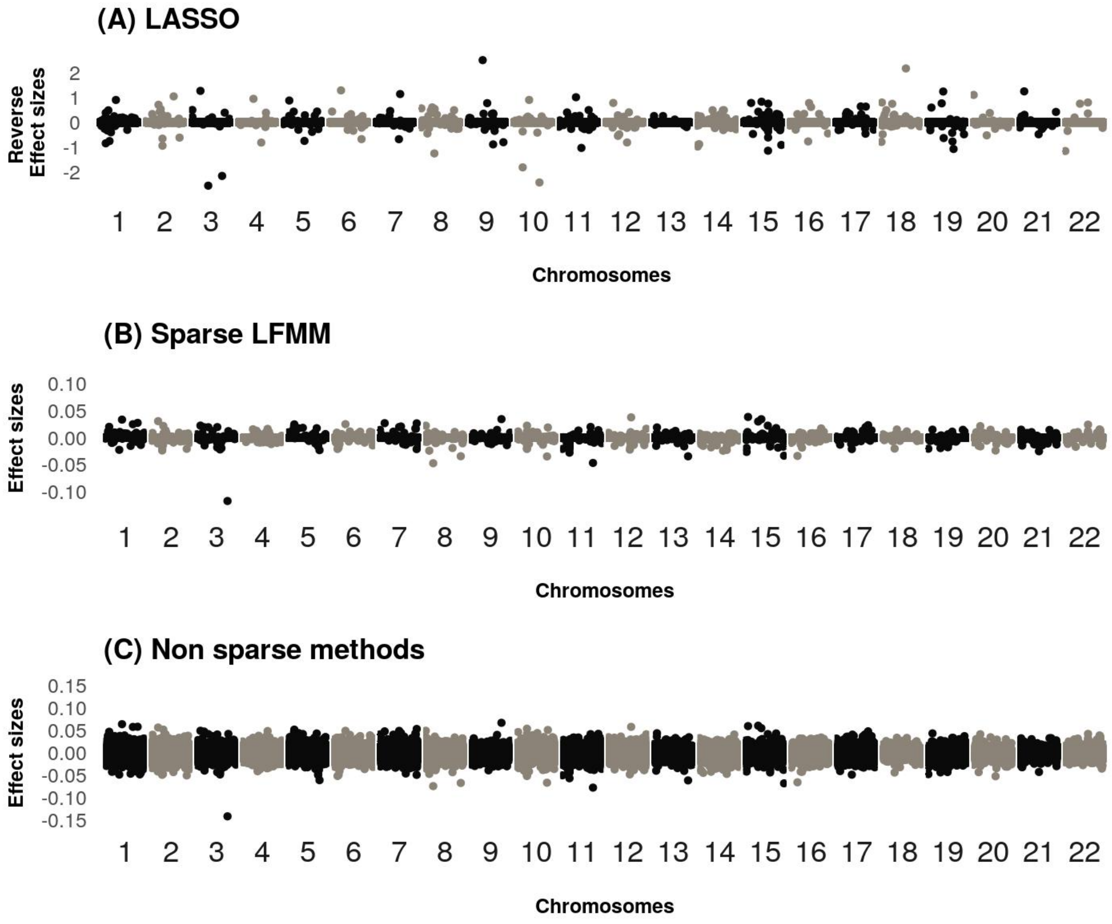


**Figure S7. DNA methylation EWAS of smoking status in pregnant women (all chromosomes).** A) Estimated reverse effect sizes for LASSO. B) Estimated effect sizes for sparse LFMM. C) Estimated effect sizes for non-sparse methods (ridge LFMM, CATE and SVA).

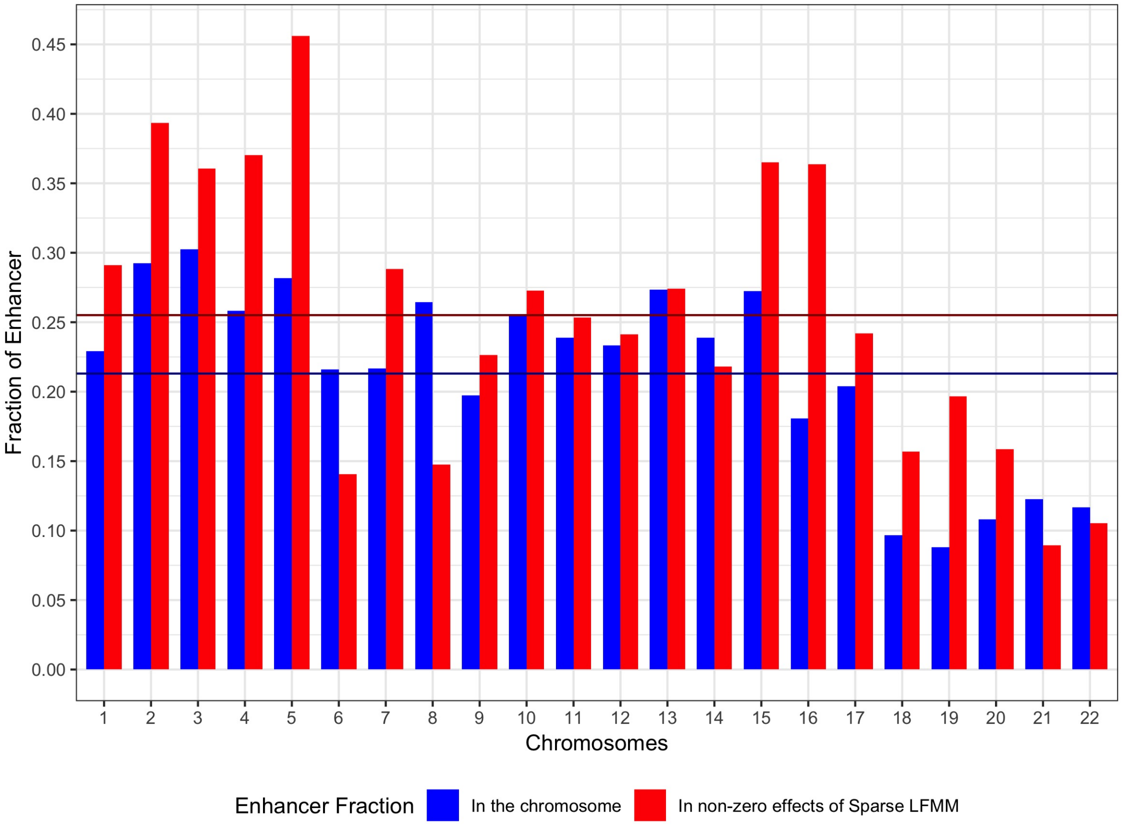


**Figure S8. EWAS of smoking status in pregnant women. Over-representation of enhancer regions in sparse LFMM CpG hits compared to the methylome.** Blue bars correspond to the fraction of enhancer regions in each chromosome. Red bars correspond to the fraction of enhancer regions detected by sparse LFMM. The horizontal blue line represent the average number of enhancer regions per chromosome for the methylome. The red line represents the average number for sparse LFMM.

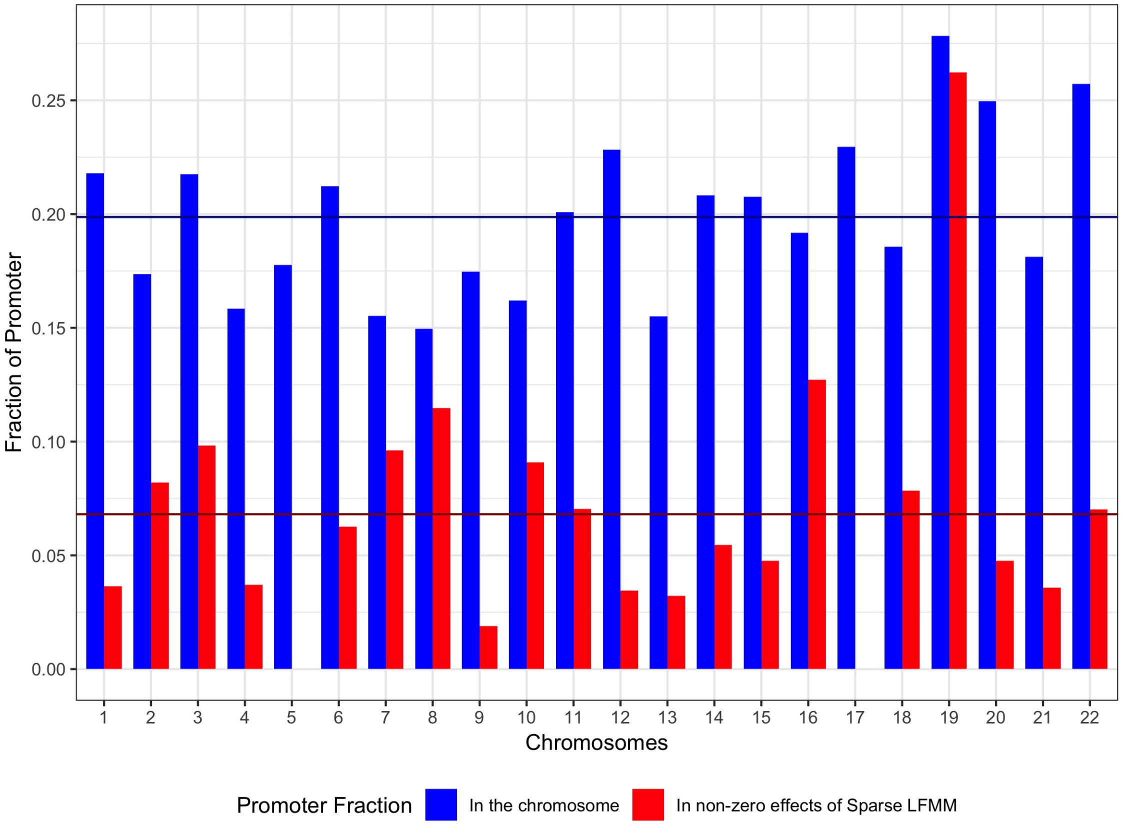


**Figure S9. EWAS of smoking status in pregnant women. Under-representation of promoter regions in sparse LFMM CpG hits compared to the methylome.** Blue bars correspond to the fraction of promoter regions in each chromosome. Red bars correspond to the fraction of promoter regions detected by sparse LFMM. The horizontal blue line represent the average number of promoter regions per chromosome for the methylome. The red line represents the average number for sparse LFMM.

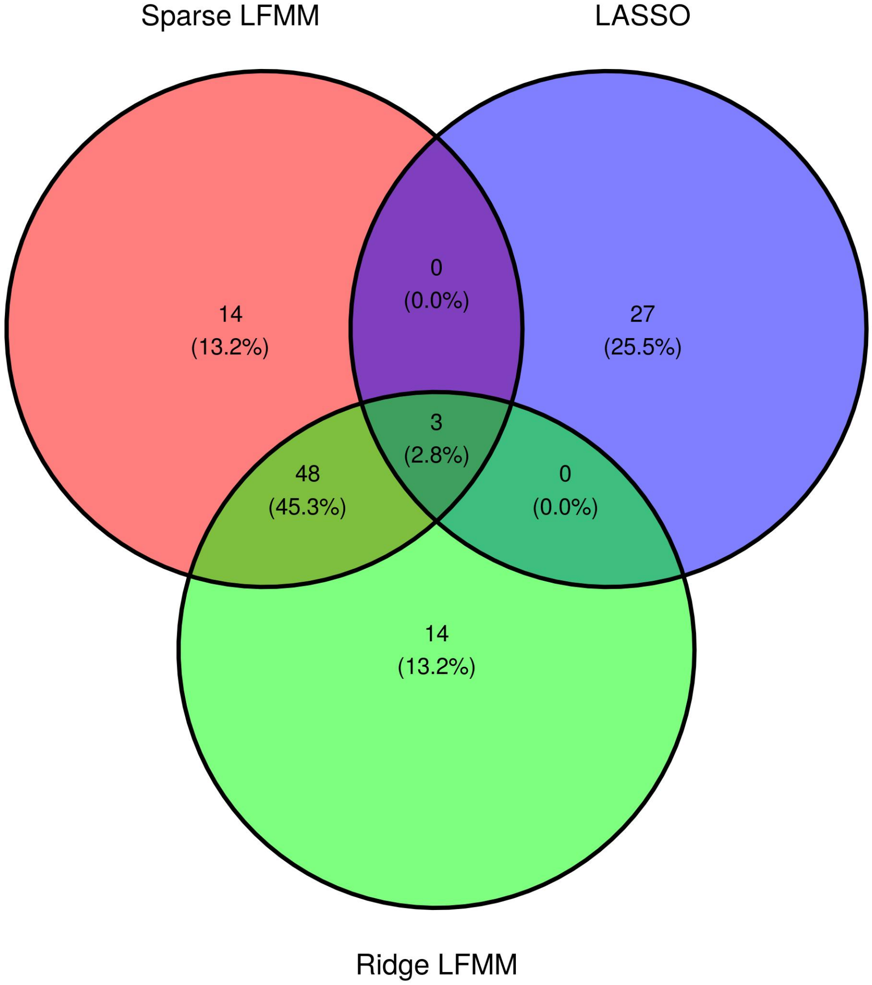


**Figure S10. EWAS of smoking status in pregnant women with sparse LFMM, LASSO and ridge LFMM**. Venn diagram of CpG hits associated with tobacco consumption in each approach. For sparse LFMM, the hits correspond to the 5% of the highest non-null effect sizes (65 hits). For LASSO, the hits correspond to the 5% of the highest non-null effect sizes (30 hits). For ridge LFMM, the hits correspond to the 65 CpGs having the highest effect sizes.

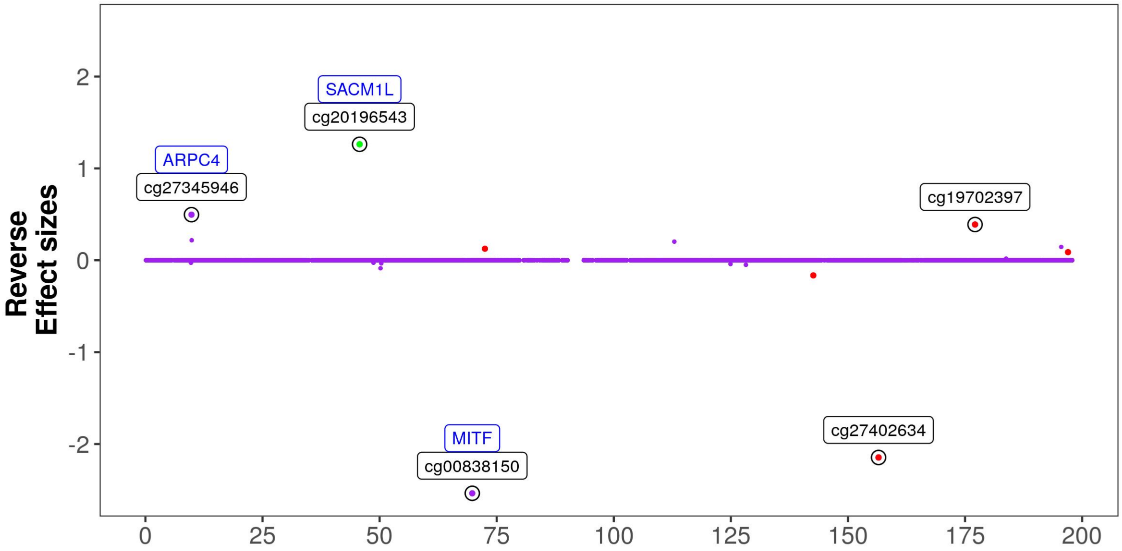


**Figure S11. Estimated reverse effect sizes for LASSO in an EWAS of smoking status in pregnant women (chromosome 3).** The 5 CpGs having the highest reverse effect sizes are circled (the genes associated with these CpGs are in blue color). Red dots represent CpGs located in enhancer regions. Green dots represent CpGs located in promoter regions (Illumina annotations).
